# IR-B deficiency and fatty acid dysregulation accelerate prostate cancer progression via PI3K/AKT signaling

**DOI:** 10.64898/2026.01.30.702723

**Authors:** Gena Huang, Athba AlQahtani, Jing Huang, Jinyu Li, Sichen Liu, Kui Jiang, Zimeng Song, Yue Xi, Shujing Wang, Man Li, Yingjie Wu

**Affiliations:** Institute for Genome Engineered Animal Models of Human Diseases, National Center of Genetically Engineered Animal Models for International Research, Liaoning Province Key Lab of Genetically Engineered Animal Models, Dalian Medical University, Dalian 116044, China; Department of Gynecological and urogenital oncology department, Department of Breast Oncology, The Second Hospital of Dalian Medical University, Dalian, Liaoning, 116044, China; Key Laboratory of Endocrine Glucose & Lipids Metabolism and Brain Aging, Ministry of Education, Shandong Provincial Hospital; Science and Technology Innovation Center, School of Laboratory Animal & Shandong Laboratory Animal Center, Shandong First Medical University & Shandong Academy of Medical Sciences, Jinan, Shandong 250021, China; Department of Endocrinology and Metabolism, The Third Affiliated Hospital of Jinzhou Medical University, Jinzhou, Liaoning 121002, China; Department of Biochemistry and Molecular Biology, Liaoning Provincial Core Lab of Glycobiology and Glycoengineering, College of Basic Medical Sciences, Dalian Medical University, Dalian, 116044, China

**Keywords:** Insulin receptor, IR-B isoform, glucolipid metabolism, prostate cancer, omega-3 fatty acid

## Abstract

The insulin receptor (IR) is markedly overexpressed in both human and mouse prostate cancer, with a significant elevation in the IR-A/IR-B ratio across patient tissues, cell lines, and *Hi-Myc* mouse prostates. To elucidate the role of IR-B in prostate tumorigenesis, we generated a prostate-specific IR-B knockout (KO) mouse model using *Pbsn-Cre*–driven recombination. Prostate-restricted loss of IR-B was confirmed at the transcript level and did not affect other tissues. Crossing these mice with *Hi-Myc* transgenics revealed that IR-B deficiency promotes accelerated progression to invasive adenocarcinoma, characterized by enhanced cellular proliferation and atypical histopathology. Transcriptomic and metabolomic profiling of dorsolateral prostate lobes demonstrated activation of PI3K/AKT and mTOR signaling, along with upregulation of IRS1/2/4 and IGF2. Metabolite analyses indicated elevated fatty acid levels and enhanced lipolysis pathways, implicating metabolic reprogramming in tumor progression. Notably, glucose and lipid metabolism genes, including GLUT1, GLUT12, FASN, and GPR120, were upregulated, accompanied by an increased BCL2/BAX ratio, suggesting apoptosis inhibition. Functional studies further revealed opposing roles of dietary fatty acids: ω-3 polyunsaturated fatty acids (EPA, DHA) suppressed prostate cancer cell survival, proliferation, and PI3K/AKT signaling while promoting apoptosis, whereas ω-6 fatty acid (arachidonic acid) exerted pro-tumorigenic, anti-apoptotic effects. Collectively, these findings identify IR-B loss as a driver of metabolic and signaling reprogramming that accelerates prostate tumorigenesis, while highlighting ω-3 fatty acids as potential modulators counteracting IR-B-deficient prostate cancer progression.

## INTRODUCTION

Hyperinsulinemia, insulin resistance, and other metabolic disorders are closely associated with the occurrence and development of carcinomas (1, 2). The roles of the insulin-like growth factor (IGF) and insulin signaling pathways in prostate cancer initiation and progression are now well established (3, 4). The insulin receptor (IR) is frequently overexpressed in prostate tumors, exerting oncogenic effects in addition to its metabolic functions, beyond regulating metabolism, insulin receptor also promotes mitogenic activity in malignant tumors. Because IR participates in both mitogenic and metabolic pathways, direct inhibition of its pro-oncogenic signaling would likely cause serious side effects. Therefore, alternative therapeutic strategies are being actively explored.

There are two major splicing variants of IR, resulting in two isoforms IR-A and IR-B with distinctive features and expression patterns in various organs and tissues. IR-B, a product of exon 11 retention, contains extra 12 amino acids compared to IR-A, generated by exon 11 skipping. While IR-A has similarly high affinity to insulin, IGF-1 and IGF-2 ligands, IR-B and IR-A/IR-B heterodimer only bind to insulin strongly with much weaker affinity for IGF-1/2 ligands. Therefore, predominant IR-A expression may drive prenatal growth and development whereas IR-B is more essential for metabolic insulin action in adults. This splicing event is evolutionarily conserved in mammals and may represent an additional layer of regulatory control, fine-tuning the pleiotropic effects of the insulin/IGF system by adjusting the amplitude of ligand-receptor interactions (5). IR-A plays a predominant role in tumors and embryonic tissues while IR-B is more important in metabolic processes (6, 7). Dysregulation of IR-A and IR-B expression has been linked to insulin resistance, aging and tumor development, with IR-A playing a greater role in carcinomas (8, 9). Increasing evidence suggests that IR-mediated signaling can recruit IRS proteins and activate PI3K and MAPK pathways, contributing to various disorders such as insulin resistance and cancer (10). Consequently, the IR-A signaling pathway has been considered as a potential therapeutic target in malignant cancer (11, 12).

Extensive research has established that dysregulated glucose and lipids metabolism is associated with higher cancer incidences (13, 14). Prostate cancer often acquire energy from lipid metabolism, including lipid regeneration and the synthesis of unsaturated fatty acids (15). Polyunsaturated fatty acids (PUFA) play important roles in prostate cancer. While the functions of omega-6 (ω-6) PUFA are well characterized, the reported roles of omega-3 (ω-3) PUFA in cancer remain contradictory (16, 17). GPR120, a known receptor for ω-3 fatty acids, has been linked to metabolic disorders as well as the growth, progress, prognosis and survival outcomes of various tumors (18–20).

To date, no antibody has been able to reliably distinguish IR-A from IR-B in conventional immunohistochemistry (IHC). Consequently, studies investigating the roles and differences of IR isoforms have primarily relied on mRNA detection. Furthermore, the lack of an IR isoform-specific knockout animal model has limited precise mechanistic exploration. In this study, we examined the IR-A/IR-B ratio in human and mouse prostate cancer. We also established a prostate-specific IR-B knockout *Hi-Myc* mouse model and demonstrated that IR-B loss acts as a driver of prostate carcinoma *in vivo*.

## RESULTS

### Insulin receptor is highly overexpressed in human and mouse prostate cancer

We analyzed the alteration frequencies of the insulin receptor *INSR* gene across four histopathologic subtypes of prostate cancers retrieved from the cBioPortal. Recurrent *INSR* gene amplification was frequently observed in more aggressive forms, such as prostate neuroendocrine carcinoma and castration-resistant prostate cancer (Figure 1A), whereas other alteration types-such as mutation or deletion of INSR-were rare. Consistent with these findings, analysis of Oncomine gene expression datasets (21, 22) revealed that *INSR* is highly overexpressed in benign prostatic hyperplasia and prostate carcinoma (Figure 1B, C). This was further supported by IHC staining, which confirmed elevated IR protein expression in prostate carcinoma tissues (Figure 1D).

**Figure 1.**
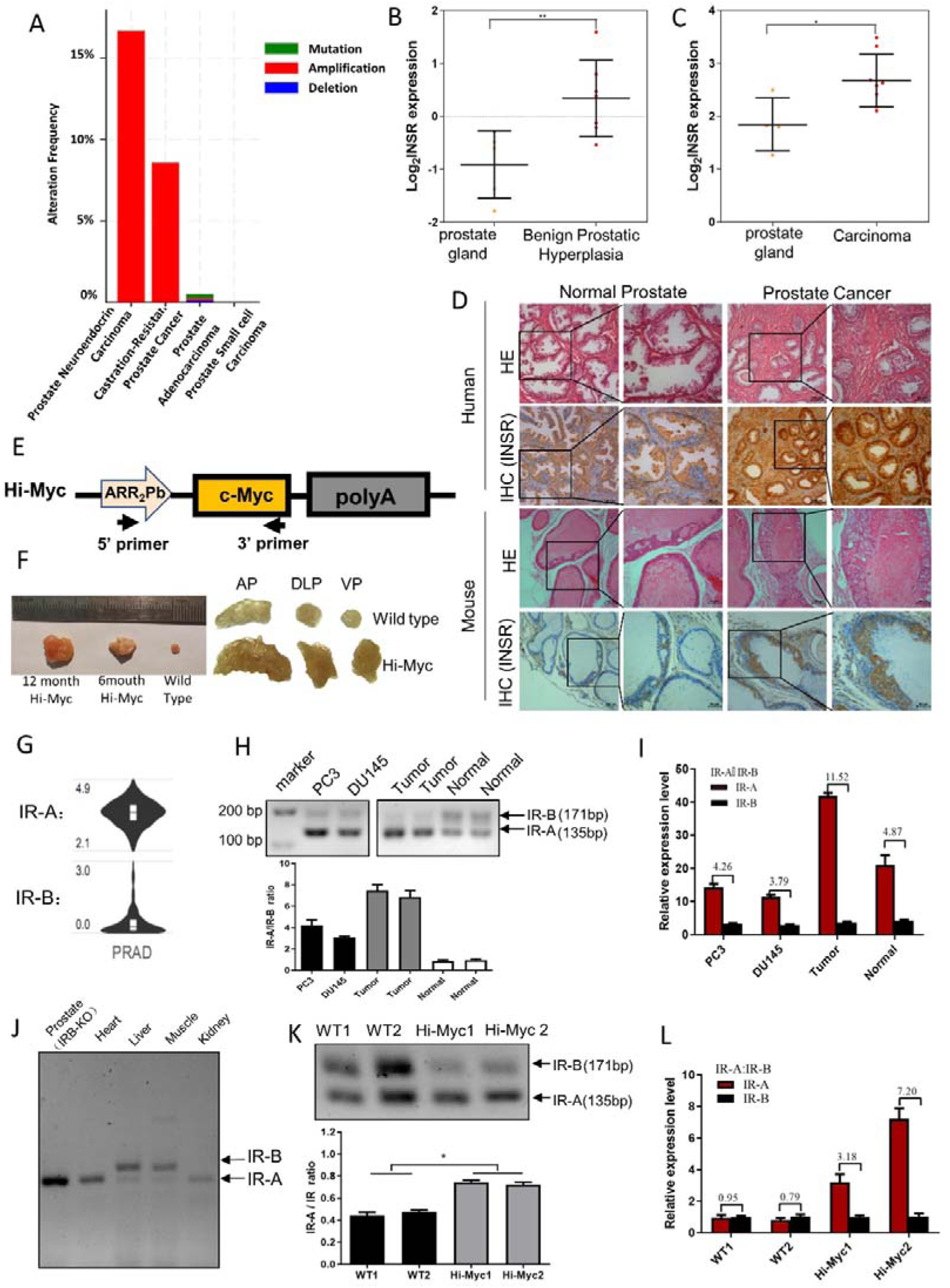
Insulin receptor and its alternative splicing isoform IR-A is highly overexpressed in prostate cancer. (A) INSR gene amplification is recurrent in prostate neuroendocrine carcinoma and castration-resistant prostate cancer based on cBioPortal data. (B) INSR expression is significantly higher in benign prostatic hyperplasia (n = 7, red circles) than in normal prostate tissue (n = 5, yellow circles), from Oncomine database. (C) INSR expression is significantly elevated in prostate carcinoma (n = 8, red circles) compared to normal prostate (n = 4, yellow circles), from Oncomine database. Black lines indicate median with interquartile range. \*\**P* < 0.01, \**P* < 0.05. (D) INSR protein levels in human and mouse prostate cancer and adjacent non-cancerous tissues assessed by IHC. Scale bar, 100 or 50 μm. (E) Schematic of *Hi-Myc* transgenic mice construction: human *c-Myc* cDNA under probasin (Pb) promoter. (F) Anatomy overview of prostate lobes in wild-type and *Hi-Myc* mice at 6 and 12 months. AP: anterior prostate; VP: ventral prostate; DLP: dorsal-lateral prostate. (G) IR-A and IR-B expression in prostate cancer from isoform usage profiling in Gene Expression Profiling Interactive Analysis (GEPIA) database. (H & I) IR splicing analyzed by Rt–PCR (H) and qPCR (I) in PC3, DU145 cells, clinical prostate cancer, and normal tissues. IR-A/IR-B ratios shown in (I). Data are mean ± SEM. (J) IR-A and IR-B mRNA expression in prostate DLP lobes and other organs of IR-B KO mice measured by PCR spanning exons 10-12. (K & L) IR splicing analysis in wild type and *Hi-Myc* mouse prostate tissues by RT–PCR (K) and qPCR (L). IR-A/IR-B ratios quantified and indicated. Data are mean ± SEM. \**P* < 0.05.

To examine IR expression in mouse prostate carcinoma tissues, we used the *Hi-Myc* transgenic model developed by Sawyers’s lab, in which human *C-MYC* is overexpressed in the prostate under the *ARR2Pb* promoter (23) (Figure 1E). This model develops prostatic intraepithelial neoplasia (PIN) as early as 2 weeks of age, progressing to high grade PIN (HG-PIN) or invasive adenocarcinoma between 6 to 12 months. The mouse prostate comprises three distinct lobes-ventral, dorsolateral, and anterior (Supplementary Figure S2A)(24)-each with unique histologic and physiologic characteristics. The dorsolateral prostate lobes (DLP), which are most analogous to the peripheral zone, are commonly used to model human prostate cancer. Thus, we performed prostate microdissection on 6-month-old wild-type and *Hi-Myc* mice. Compared with wild-type mice, *Hi-Myc* prostates-and their individual lobes-were larger, firmer, and exhibited altered texture (Figure 1F, Supplementary Figure S2B). IHC staining confirmed that, consistent with human data, IR protein was also overexpressed in Hi-Myc mouse prostate cancer tissues (Figure 1D).

We next investigated whether IR isoforms were differentially expressed in various caner types. Using isoform usage profiling in GEPIA, we found that IR-A was highly expressed across multiple cancer types, including prostate cancer, and was associated with poor prognosis (Figure 1G, Supplementary Figure S1). Differential expression analysis further revealed that the IR-A/IR-B ratio was significantly elevated in prostate cancer cell lines, patient tissues, and Hi-Myc mouse prostates compared with their respective controls (Figure 1H-I, 1K-L). Together, these findings demonstrate that both *INSR* expression and the IR-A/IR-B ratio are markedly increased in human and mouse prostate cancer.

### Establishment of a mouse model for IR-B-deficient prostate cancer

We previously reported that IR-A is the predominant isoform in breast cancer and that IR-B knockout accelerates breast cancer progression. However, the lack of an *in vivo* IR-B knockout model has limited similar investigations in prostate cancer. To address this, we generated a novel prostate-specific IR-B knockout mouse model. LoxP-exon11-loxP (IR-B^Floxed^) mice were crossed with Pbsn-Cre mice, producing offspring with prostate-specific deletion of IR-B (Figure 2A-C). Efficient Pbsn-Cre recombination in prostate tissues was confirmed by crossing Pbsn-Cre mice with floxed tdTomato reporter mice (Figure 2D). tdTomato fluorescence was detected in the prostates of Pbsn-Cre/tdTomato mice but absent in tdTomato-only controls, confirming the specificity of Pbsn promoter activity. Among the three prostate lobes, fluorescence intensity was highest in the anterior prostate (AP) and dorsolateral prostate (DLP) lobes, and lower in the ventral prostate (VP) (Figure 2E). Importantly, Cre activity driven by the Pbsn promoter induced prostate-specific IR-B disruption without affecting other organs or tissues (Figure 1J). Loss of IR-B in the prostate was further validated at the mRNA level, where IR-B transcripts were significantly reduced in the DLP lobes of IR-B knockout mice (Figures 2F-H). These results confirm the successful establishment of a prostate-specific IR-B deficient mouse model, enabling *in vivo* investigation of IR-B function in prostate cancer.

**Figure 2:**
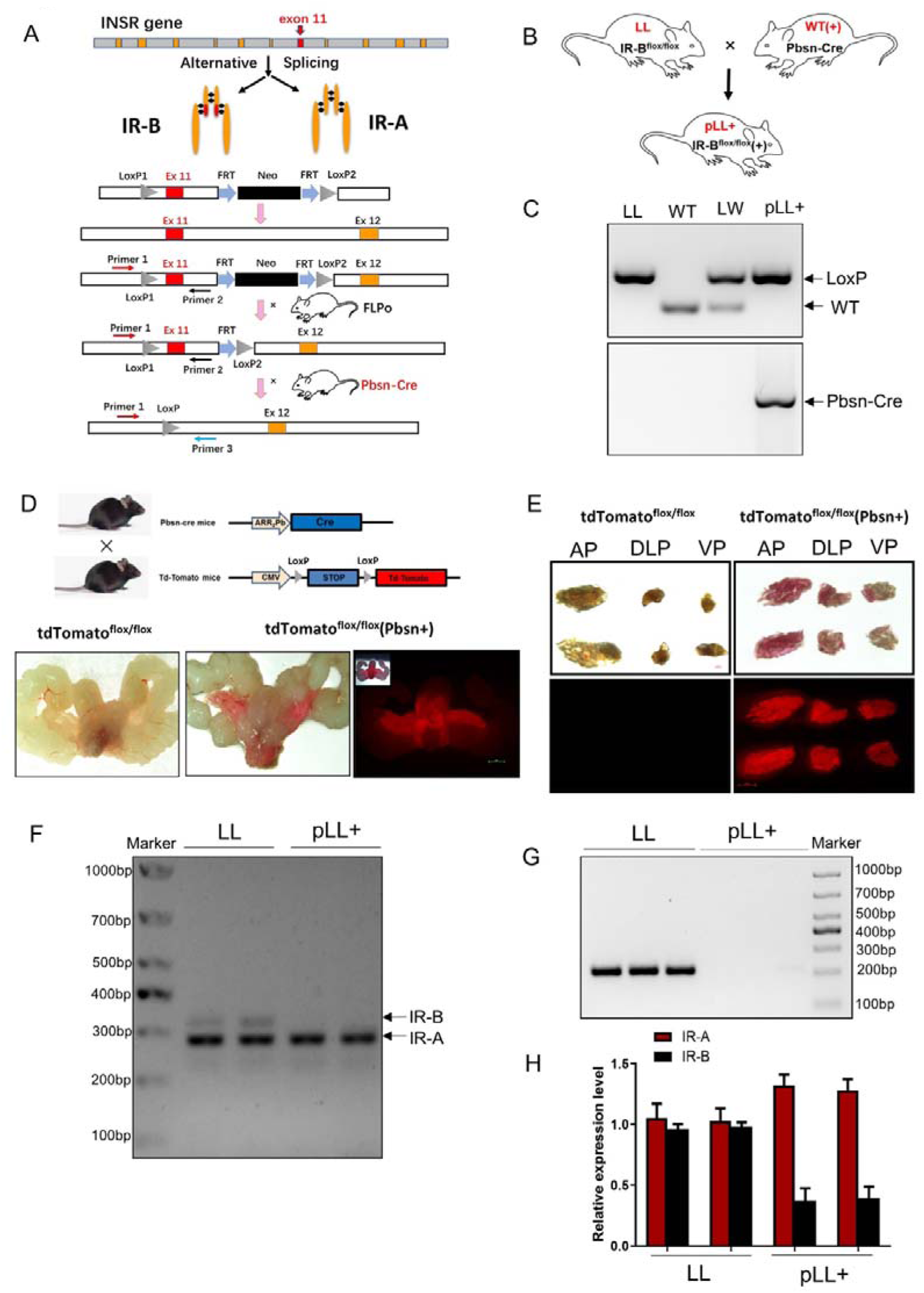
Construction of prostate-specific IR-B knockout mice. (A & B) Schematic diagram showing the process of generating prostate-specific IR-B KO mice used in this study, where WT refers to wild-type, LL to LoxP sites in floxed IR-B mice, pLL to IR-B KO mice, and (+) indicates the presence of Pbsn-Cre transgene. (C) PCR genotyping of LoxP sites in floxed IR-B mice (LL) and IR-B KO mice (pLL^+^). Pbsn-Cre transgene only detected in IR-B KO mice (pLL^+^). (D) Pbsn-Cre activity confirmed with floxed tdTomato mice to remove the STOP signal and allow tdTomato protein expression. (E) Microscopic and immunofluorescence of prostate lobes in floxed tdTomato and tdTomato^flox/flox^ (Pbsn+) mice; AP: anterior prostate; VP: ventral prostate; DLP: dorsal and lateral prostate. (F & G) IR-A and IR-B gene expression in the prostate DLP lobes of floxed IR-B mice (LL) and IR-B KO mice (pLL^+^), with PCR primers spanning exons 10 and 12 (F) and PCR primers spanning exons 10 and 11 (G). (H) Relative gene expression of IR-A and IR-B in prostate DLP lobes of LL mice and pLL^+^ mice measured by real-time PCR. Data was expressed as means ± SEM.

### Prostate growth and development are independent of IR-B

IR-B floxed (LL) and prostate-specific IR-B knockout (pLL^+^) mice were monitored from 3 to 6 months of age. No significant differences were observed in body weight, prostate lobe volume, or tissue texture between the two groups (Figures 3A-E). Histological examination of DLP lobes by H&E staining revealed normal morphology in both LL and pLL^+^ mice at 3 and 6months. The incidence of prostatic intraepithelial neoplasia (PIN) was comparable between the groups at both time points (Figures 3F-G). These results indicate that IR-B is not essential for normal prostate growth or baseline histological development.

**Figure 3.**
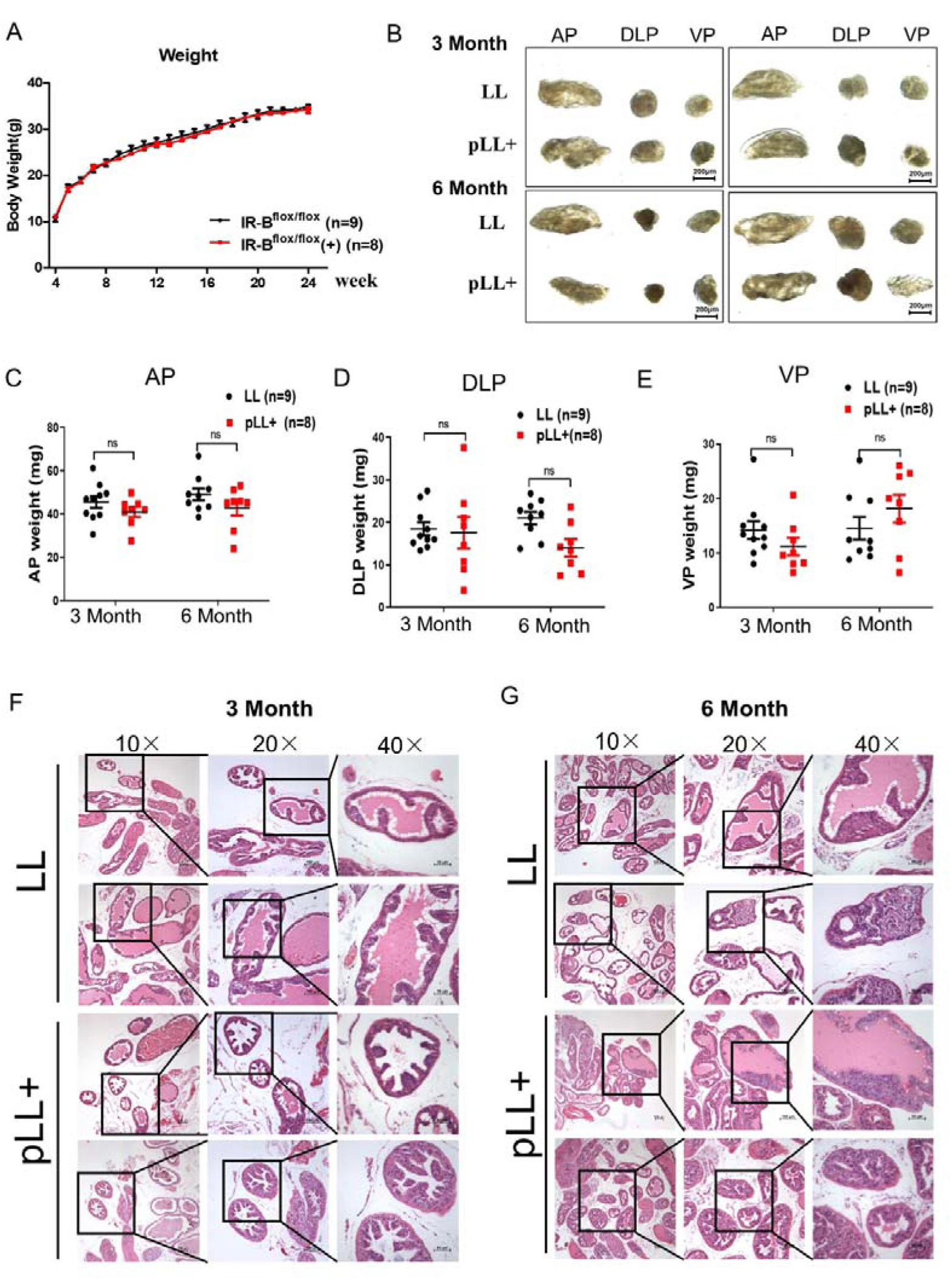
Prostate IR-B knockout has no significant effect on the size and development of prostate gland. (A) Body weight of IR-B ^flox/flox^ (LL) mice (n=9) and IR-B ^flox/flox^(+) (pLL^+^) mice (n=8) at various age. (B) Representative prostate lobe images of LL and pLL^+^ mice at 3 and 6 months. Scale bar, 200 μm. (C & E) Weights of AP (C), DLP (D) and VP (E) lobes in LL (n=9) and pLL^+^ (n=8) mice at 3 and 6 months. Data are mean ± SEM, ns = not significant. (F & G) H&E staining of DLP lobes showing histology of LL and pLL^+^ mice at 3 (F) and 6 months (G). Scale bar =150, 100, 50 μm (10X, 20X and 40X).

### Prostate IR-B knockout promotes more aggressive prostate adenocarcinoma in *Hi-Myc* transgenic mice

Considering that insulin receptor isoforms are often linked to cell proliferation in neoplastic tissues, investigating their role in prostate carcinoma is essential. To assess the impact of IR-B knockout on prostate cancer development and progression, we crossed *Hi-Myc* transgenic mice with IR-B floxed mice, generating prostate-specific IR-B knockout *Hi-Myc* mice (pLL^+^H) (Supplementary Figure S3).

Hi-Myc mice are known to develop PIN as early as 2 weeks of age, progressing to invasive adenocarcinoma by 6 to 12 months (23). Therefore, we conducted microdissection of prostates at 3 and 6 months. Significant differences in appearance and weight of prostate lobes were observed between IR-B floxed *Hi-Myc* mice (LL^H^) and IR-B prostate knockout *Hi-Myc* mice (pLL^+H^) at 3 months (Figures 4A-E), with these differences becoming more pronounced at 6 months (Figures 5A-D). Histological analysis of the DLP using H&E staining revealed that, compared to LL^H^ and pLL^+H^ mice exhibited a higher incidence of PIN at 3 months (Figure 4F) and a significantly increased incidence of invasive prostate cancer at 6 months (Figure 5E, F). Cytological atypia characterized by enlarged nuclei (Figure 4G; Figure 5G, H) and elevated Ki-67 positive cell counts were observed in pLL^+H^ mice at both time points (Figure 4H, Figures 5I-K). Additionally, aberrant androgen receptor (AR) expression and reduced apoptosis, assessed by IHC and immunofluorescence, were detected in 6-month pLL^+H^mice (Supplementary Figure S4).

**Figure 4.**
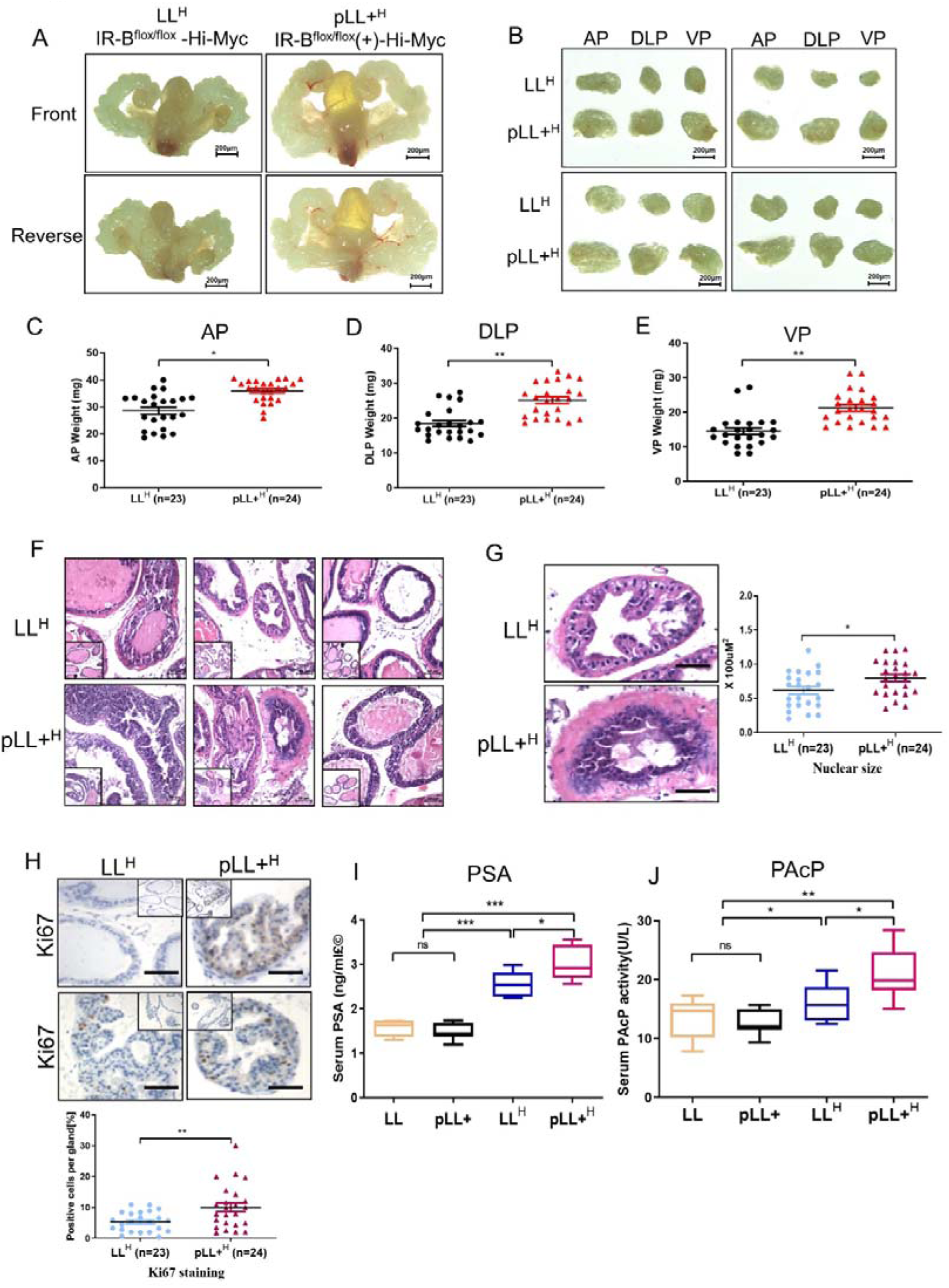
Prostate IR-B knockout in *Hi-Myc* mice leads to severe prostatic intraepithelial neoplasia. (A) Images of intact prostate glands from IR-B^flox/flox^-*Hi-Myc* (LL^H^) and IR-B^flox/flox^(+)-*Hi-Myc* mice (pLL^+H^) at 3 months. front and back views shown. Scale bar, 200 μm. (B) Representative images of prostate lobes from LL^H^ and pLL^+H^ mice at 3 months. Scale bar, 200 μm. (C, D & E) Weights of AP (C), DLP (D) and VP (E) in LL^H^ (n=23) and pLL^+H^ (n=24) mice at 3 months. Data mean ± SEM. \*\**P* < 0.01, \**P* < 0.05. (F) H&E staining of DLP lobes at 3 months showing histological differences between LL^H^ and pLL^+H^. Scale bar, 50 μm. (G) Left: Representative nuclei on H&E sections; Right: Quantification of nuclear size (largest 20 nuclei) in LL^H^ (n = 23) and pLL^+H^ mice (n = 24). Scale bar, 100 μm. \**P* < 0.05. (H) Ki-67 IHC in DLP lobes of 3-month-old LL^H^ and pLL^+H^ mice; Quantification shown. Scale bar, 100 μm. \*\**P* < 0.01. (I & J) Serum PSA (I) and PAcP (J) levels in LL (n=9), pLL^+^ (n=8), LL^H^ (n=23) and pLL^+H^ mice (n=24) at 3 months. ns = not significant. \*\*\**P* < 0.001, \*\**P* < 0.01, \**P* < 0.05.

**Figure 5.**
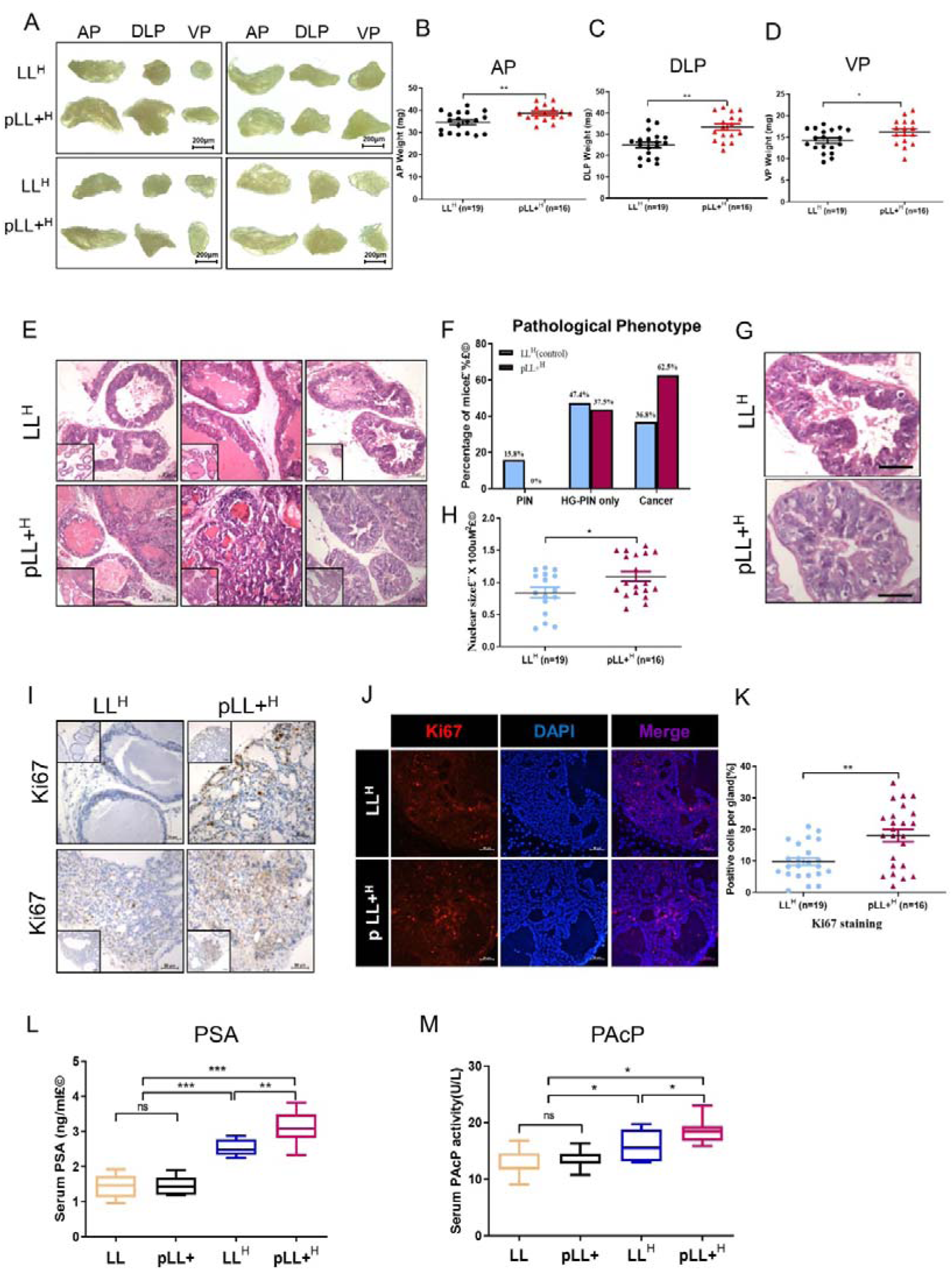
Prostate IR-B knockout in *Hi-Myc* transgenic mice leads to invasive adenocarcinoma. (A) Representative prostate lobes images from IR-B^flox/flox^-*Hi-Myc* (LL^H^) and IR-B^flox/flox^(+)-*Hi-Myc* mice (pLL^+H^) at 6 months. Scale bar, 200 μm. (B, C & D) Weights of AP (B), DLP (C) and VP (D) lobes in LL^H^ (n=19) and pLL^+H^ (n=16) mice at 6 months. Data mean ± SEM. \*\**P* < 0.01, \**P* < 0.05. (E) H&E staining of DLP lobes at 6 months. Scale bar, 50 μm. (F) Incidence of PIN, high-grade PIN (HG-PIN), and invasive cancer pLL^+H^ and LL^H^ mice. (G) Representative nuclei in H&E sections from LL^H^ and pLL^+H^ mice. Scale bar, 100 μm. (H) Nuclear size quantification (largest 20 nuclei) at 6 months. \**P* < 0.05. (I, J & K) Ki-67 staining by IHC and immunofluorescence in DLP lobes; quantification shown. Scale bar, 50 μm. \*\**P* < 0.01. (L & M) Serum PSA (L) and PAcP activity (M) in LL (n=9), pLL^+^ (n=8), LL^H^ (n=19) and pLL^+H^ (n=16) mice at 6 months. Data mean ± SEM. ns = not significant; \*\*\**P* < 0.001, \*\**P* < 0.01, \**P* < 0.05.

Serum prostate-specific antigen (PSA) and prostate acid phosphatase (PAcP) are effective markers for distinguishing early prostate carcinoma or prostatitis from benign prostate hyperplasia (29). We measured serum PSA levels and PAcP activity at 3 and 6 months, finding no difference between the LL and pLL^+^ groups. However, both LL^H^ and pLL^+H^ mice showed significantly elevated PSA and PAcP activity, with higher levels in pLL+^H^ mice (Figure 4I, J; Figure 5L, M).

Together, these findings suggest that prostate-specific deletion of IR-B in *Hi-Myc* mice promotes cellular proliferation with atypical features and accelerates progression toward invasive carcinoma, highlighting the role of IR-B in prostate cancer development within this genetic context.

### IR-B KO/*Hi-Myc* mice exhibit elevated activation of PI3K/AKT signaling

To further explore signaling pathways affected by IR-B knockout and assess the relevance of this models to human prostate cancer, we performed RNA sequencing (RNA-seq) and targeted metabolomics using UPLC-MS/MS on the prostate DLP lobes of IR-B KO/ *Hi-Myc* (pLL^+H^) and control mice (LL^H^).

RNA-seq analysis revealed significant differential expression of 539 genes (log2FC > 1, adjusted *p* < 0.05), with 262 genes upregulated and 277 downregulated in IR-B knockout prostates compared to controls (Figure 6A). Hierarchical clustering based on IR-B status highlighted enrichment of mTOR signaling and cell proliferation pathway, along with genomic features known to be associated with insulin receptor (30) (Figure 6B). KEGG pathway analysis showed that upregulated genes were primarily involved in mTOR signaling, prostate cancer, and regulation of lipolysis in adipocytes—processes that may promote tumor progression (Figure 6C, Supplementary Table 2). Consistent with these findings, key components of the PI3K/AKT pathway, including IRS-1, IRS-2, IRS-4 and IGF2, were upregulated in the IR-B knockout group (Figure 6D, E).

**Figure 6.**
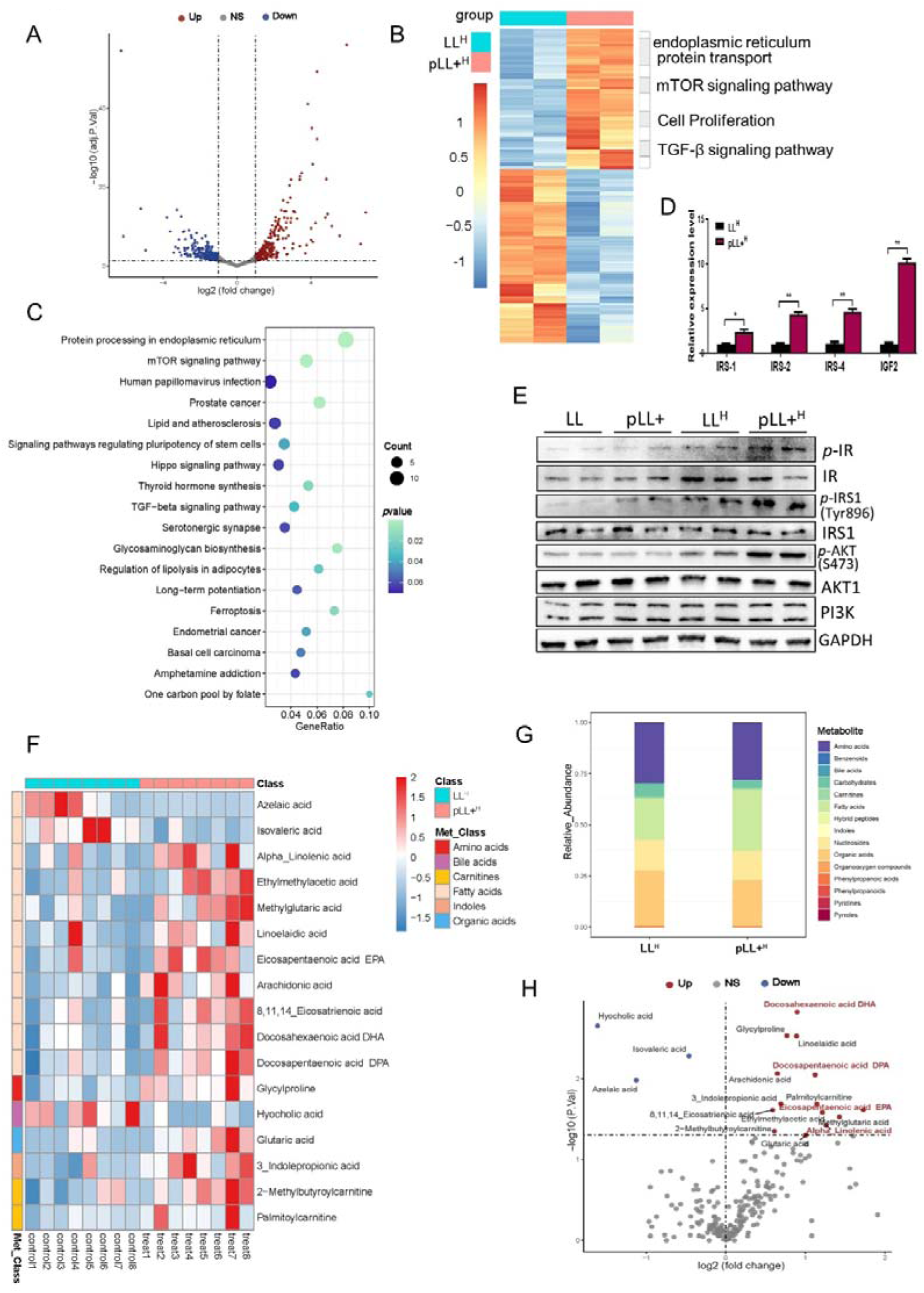
Transcriptomics and metabolomics analysis of prostate IR-B knockout in *Hi-Myc* transgenic mouse prostate gland. (A) Volcano plot showing differentially expressed genes between LL^H^ (control) and pLL^+H^ prostate DLP lobes. (B) Hierarchical clustering and heatmap of significantly differentially expressed genes; enriched gene ontology (GO) terms indicated. (C) Bubble plot of enriched KEGG pathways for upregulated genes. (D) Real-time PCR quantification of IRS-1, IRS-2, IRS-3, and IGF2 mRNA in LL^H^ and pLL^+H^ prostate DLP lobes. mean ± SEM. (E) Western blot analysis of *p-*IR, total IR, *p-*IRS, total IRS1, *p-*AKT, total AKT, and PI3K protein in LL, pLL^+^, LL^H^ and pLL^+H^ prostate DLP lobes. (F) Heatmap showing differential metabolites between pLL^+H^ and LL^H^ mice. (G) Classification of metabolite type in prostate DLP lobes of LL^H^ and pLL^+H^ mice. (H) Volcano plot of significantly differential metabolites; fatty acids highlighted.

To identify metabolic alterations induced by IR-B knockout, targeted metabolomics of prostate DLP lobes from LL^H^ and pLL^+H^ mice was conducted using ultra-performance liquid chromatography coupled with tandem mass spectrometry (UPLC-MS/MS). The analysis revealed increased levels of fatty acid in the IR-B knockout group, as demonstrated by metabolite classes distribution and Z-Score heatmaps (Figure 6F, G). Volcano plot analysis identified 14 upregulated and 3 downregulated metabolite classes (Figure 6H, Supplementary Table 3). Notably, nine of the upregulated metabolites were fatty acids, predominantly polyunsaturated fatty acids (PUFAs), including four omega-3 polyunsaturated fatty acids: docosahexaenoic acid (DHA), eicosapentaenoic acid (EPA), docosapentaenoic acid (DPA), and alpha-linolenic acid (Figure 6H). These metabolomic results align with transcriptomic data indicating activation of pathways involved in adipocytes lipolysis.

### IR-B KO/*Hi-Myc* mice exhibit glucose metabolism dysregulation

It is well established that many aggressive tumors exhibit metabolic dysregulation, with glycolysis and fatty acid synthesis closely linked to tumor vitality (31, 32). Our metabolomics analysis indicated that omega-3 polyunsaturated fatty acids were the predominant upregulated metabolites. Consequently, we focused on genes involved in fatty acid synthesis. As shown in Figure 7A, expression of GPR120 (an omega-3 fatty acid receptor) and fatty acid synthase (FASN) were significantly increased in IR-B KO/*Hi-Myc* mice. Similarly, genes related to glucose uptake and glycogen synthesis, such as GLUT1 and GLUT12, were also upregulated (Figure 7A). Additionally, the ratio of Bcl-2 to Bax—key regulators of apoptosis— was significantly elevated in prostate tissue within the *Hi-Myc* background, especially in the IR-B knockout group (Figure 7E), suggesting inhibition of apoptosis in tumor tissues.

**Figure 7.**
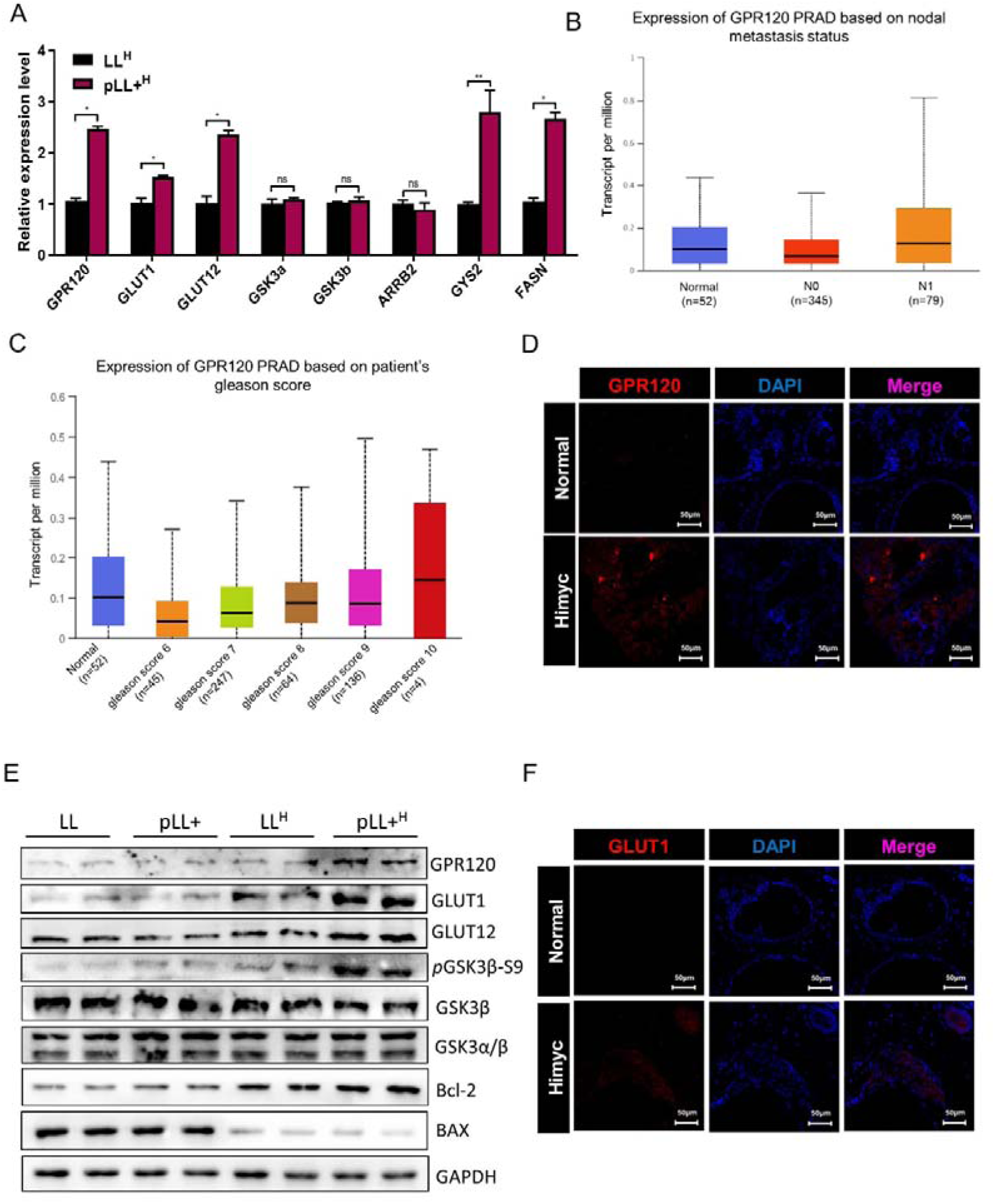
Activation of ω-3 polyunsaturated fatty acid metabolism signaling pathway by prostate IR-B knockout in *Hi-Myc* mice. (A) Relative mRNA expression of lipid synthesis-related genes in DLP lobes of LL^H^ and pLL^+H^ mice by qPCR. (B) GPR120 expression in prostate adenocarcinoma (PRAD) stratified by nodal metastasis status, from TCGA data. (C) GPR120 expression based on Gleason score in PRAD patients, TCGA data. (D) Immunofluorescence staining of GPR120 in DLP lobes of *Hi-Myc* and wild-type mice. Scale bar: 50μm. (E) Western blot of GPR120, GLUT1, GLUT12, *p*GSK3β-S9, *GSK3*β, GSK3α/β, Bcl-2, and BAX in DLP lobes of LL, pLL^+^, LL^H^ and pLL^+H^ mice. (F) Immunofluorescence of GLUT1 in DLP lobes. Scale bar: 50μm.

Since GPR120 promotes adipogenesis, we analyzed data from the TCGA database and found no significant difference in GPR120 expression between normal prostate tissue and primary prostate carcinomas (Supplementary Figure S5A, B). However, GPR120 expression was markedly increased in metastatic lesions, both in lymph nodes and distant sites, compared to normal tissue and carcinoma in situ (Figure 7B, Supplementary Figure S5C).

Furthermore, GPR120 expression correlated positively with Gleason score, rising with increasing tumor grade (Figure□7C). Consistent with these observations, both GPR120 and GLUT1 were upregulated in highly malignant prostate cancer tissues from *Hi-Myc* transgenic mice (Figure□7D,□F). Collectively, these findings underscore the critical involvement of glucose and lipid metabolic pathways in prostate carcinoma progression.

### Omega-3 (**ω**-3) fatty acids reduce cell survival and migration, while omega-6 (**ω**-6) promotes these processes

Cell viability assays demonstrated that treatment with ω-3 fatty acids (EPA and DHA) caused a time-dependent decrease in cell survival compared to controls, whereas treatment with the ω-6 fatty acid arachidonic acid (AA) resulted in a slight increase in survival over 48 hours (Figure 8A). Histological analysis corroborated these findings, showing that tissues treated with ω-3 fatty acids exhibited marked structural disorganization and follicular degeneration, while ω-6-treated tissues retained a more compact and organized architecture (Figure 8B). Functional migration assays further revealed that EPA and DHA significantly inhibited cell migration, whereas AA markedly enhanced migratory capacity compared to controls (Figure 8C, D). Additionally, ω-6 treatment was associated with a time-dependent increase in Ki67 expression, indicating elevated proliferative activity in prostate tissue, whereas ω-3 treatment suppressed Ki67 expression, suggesting reduced proliferation relative to both control and ω-6 groups (Figure 8E).

**Figure 8.**
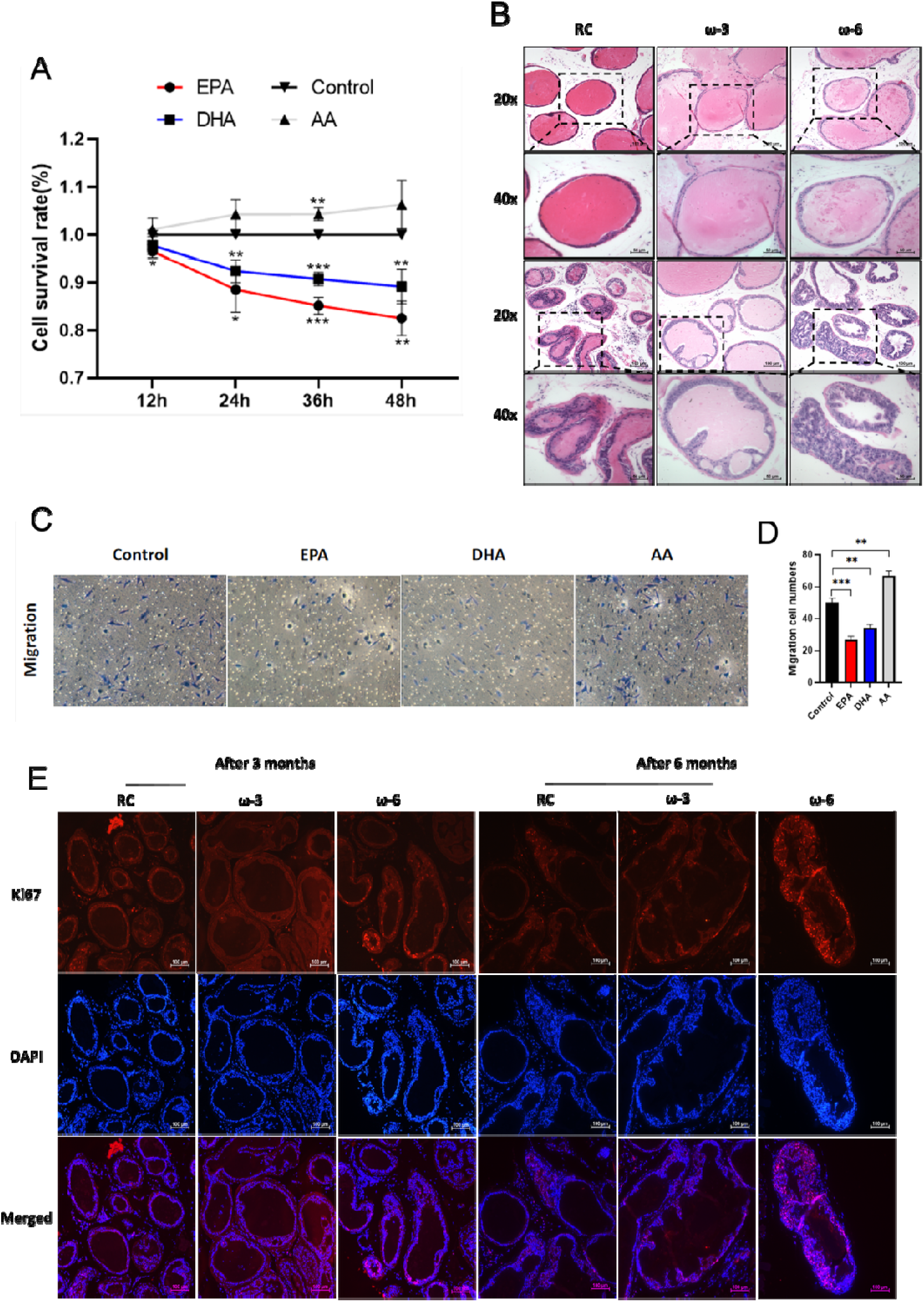
omega-3 (ω-3) fatty acids reduce cell survival and migration, while omega-6 (ω-6) promotes survival and migration. (A) Cell viability at 12, 24, 36, and 48 h after treatment with EPA (ω-3), DHA (ω-3), or AA (ω-6). EPA and DHA reduce survival; AA slightly increases survival. (B) Representative H&E images of prostate tissues from control (RC), ω-3, and ω-6 groups, showing structural disorganization with ω-3 and preserved structure with ω-6. (C & D) Cell migration assays: images and quantification showing EPA and DHA inhibit migration, AA enhances migration. Data mean ± SEM. *p<0.05, **p<0.01, ***p<0.001. (E) Immunofluorescence for Ki67 (red) and nuclei (DAPI, blue) in prostate tissues at 3□ and 6□months. ω-6 increases proliferation; ω-3 decreases proliferation compared to control.

These results support the conclusion that ω-6 fatty acids promote, while ω-3 fatty acids inhibit, epithelial cell proliferation and migration during prostate tumorigenesis. ω**-3 downregulates AR and suppresses PI3K/AKT signaling, while** ω**-6 exerts the opposite effect**

To elucidate the molecular mechanisms underlying these effects, we examined androgen receptor (AR) expression and activation of the PI3K/AKT pathway. Western blot analysis revealed a pronounced reduction in AR protein levels in the ω-3-treated groups, whereas ω-6-treatment significantly increased AR expression (Figure 9A, B). Additionally, phosphorylation of PI3K and AKT was markedly decreased in the ω-3 groups at both 3 and 6 months, whereas ω-6 treatment enhanced phosphorylation of these proteins, indicating activation of the PI3K/AKT signaling pathway (Figure 9C, D).

**Figure 9.**
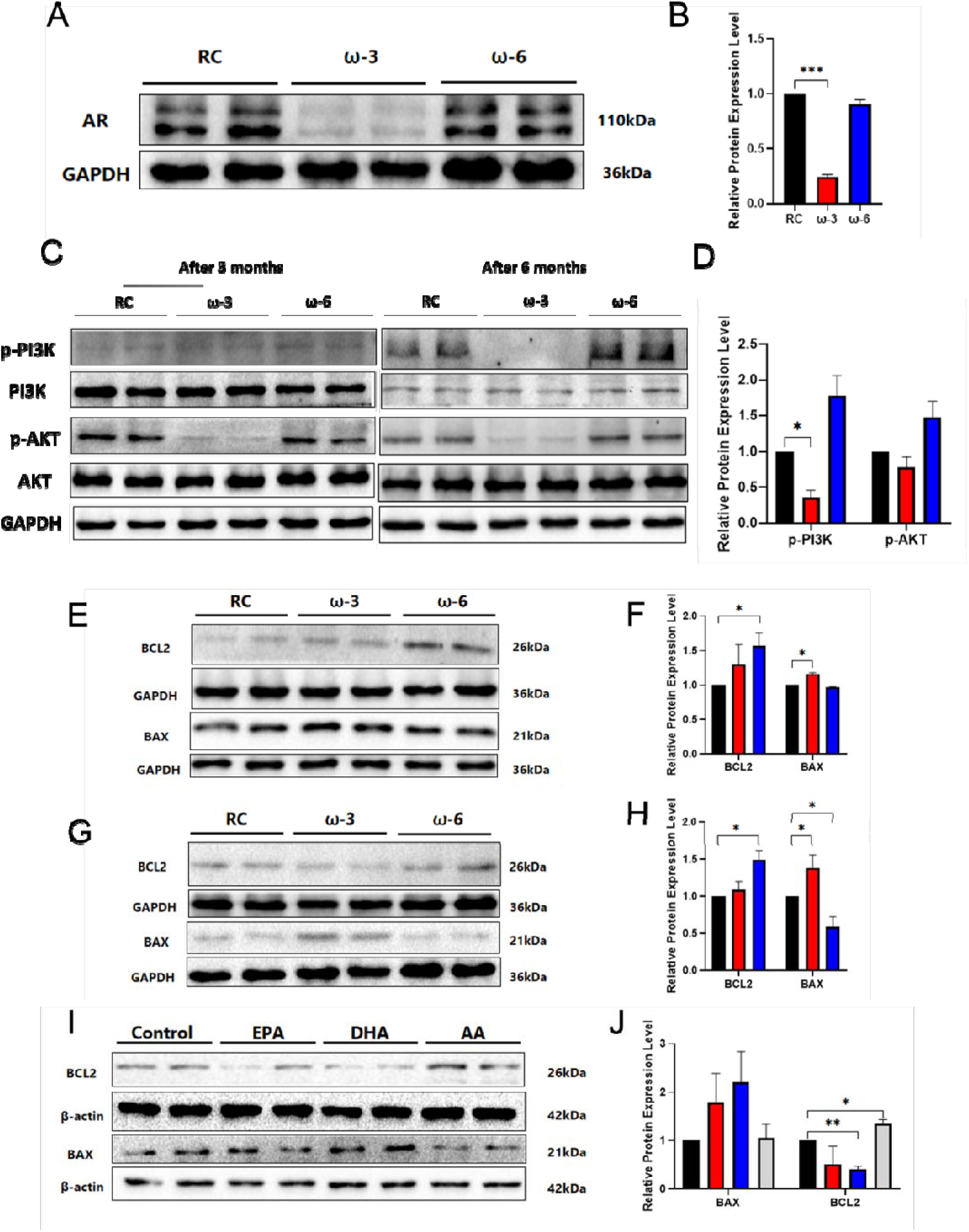
Omega-3 (ω-3) and omega-6 (ω-6) fatty acids differentially regulate AR, PI3K/AKT signaling, and apoptosis-related proteins. (A & B) Western blot and quantification showing AR expression reduced by ω-3 and increased by ω-6. (C & D) Western blot and quantification of *p*-PI3K, PI3K, *p*-AKT, and AKT at 3 and 6 months; ω-3 decreases phosphorylation, ω-6 increases it. (E, F, G, & H) Western blot and quantification of BCL2 and BAX at 3 and 6 months. ω-3 increases pro-apoptotic BAX and decreases anti-apoptotic BCL2, ω-6 shows opposite pattern. (I & J) Western blot of cultured cells treated with EPA, DHA, or AA, corfirming protein expression treads. Data mean ± SEM. *p<0.05, **p<0.01, ***p<0.001.

### **ω**-3 promotes apoptosis through BCL2/BAX modulation, whereas **ω**-6 is anti-apoptotic

Analysis of apoptosis-related proteins showed that ω-3 treatment decreased the anti-apoptotic protein BCL2 and increased the pro-apoptotic protein BAX, resulting in an elevated BAX/BCL2 ratio at both 3 and 6 months (Figure 9E–H), consistent with the findings in Figure 7. In contrast, ω-6 exposure increased BCL2 expression and reduced BAX levels, shifting the balance toward cell survival. Parallel results were observed in cultured cells treated with EPA, DHA, or AA, where EPA and DHA enhanced BAX expression and suppressed BCL2, while AA had the opposite effect (Figure 9I, J).

Collectively, these data indicate that ω-3 fatty acids suppress AR expression, inhibit PI3K/AKT signaling, and promote apoptosis, leading to reduced cell survival and migration. Conversely, ω-6 fatty acids enhance AR signaling, activate PI3K/AKT, inhibit apoptosis, and thereby promote cell survival and migratory behavior.

## DISCUSSION

The insulin receptor (IR) mediates both metabolic and mitogenic functions. Although the physiology of IR isoforms and their roles in neoplastic tissues remain incompletely understood, decreased expression of the IR-B isoform has been linked to adverse effects (5). Metabolic dysregulation contributes significantly to the progression of various diseases, including cancer (33). In this study, we generated prostate-specific IR-B knockout mice to investigate how modulation of the IR-B isoform affects glucose and lipid metabolism in prostate cancer. Our findings demonstrate that metabolic disturbances caused by reduced IR-B expression promote prostate carcinoma development *in vivo*, establishing IR-B as a key driver in prostate cancer progression.

In many tumors, IR-A expression exceeds that of IR-B (5). Previously, we showed that the IR-A/IR-B ratio is elevated in breast cancer cell lines and clinical samples compared to normal breast tissues. Manipulation of insulin receptor isoform expression via splicing factors altered oncogenic phenotypes such as proliferation, invasion and metastasis (9). Here, we similarly observed high expression of insulin receptor and increased IR-A/IR-B ratios in human and mouse prostate cancer tissues (Figure 1). This imbalance can arise from either increased IR-A or decreased IR-B expression (34). Prior studies focused on mRNA-level differences due to a lack of isoform-specific antibody, limiting insights into their physiological roles *in vivo*. By generating prostate-specific IR-B knockout mice on a *Hi-Myc* transgenic background, we unambiguously elucidated IR-B’s role in prostate cancer progression.

The insulin receptor’s signaling pathways, notably PI3K/AKT and Ras/Raf/MEK/ERK, mediate metabolic and mitogenic effects (35, 36). IGF-2 binding to IR-A activates PI3K and Ras pathway, promoting cell migration and inhibiting apoptosis, whereas IR-B primarily binds insulin and mediates differentiation and metabolic signaling (37, 38). Our data revealed activation of mTOR signaling and cell proliferation pathways in IR-B knockout mice (Figure 6B, C), along with upregulation of IRS substrates and PI3K/AKT components (Figure 6D, E). This aligns with findings in R−/IR-A cells, where IGF-2 induces mitogenic effects via Akt activation (39, 40). These results suggest that IR-A dominance favors PI3K pathway activation and mitogenic signaling. Moreover, IR isoforms can heterodimerize with IGF1R to form hybrid receptors (HR), altering ligand specificity; thus, the relative abundance of IR isoforms and IGF-1R likely shapes downstream signaling and biological outcomes (41, 42).

Metabolic reprogramming is a hallmark of tumors, facilitating rapid proliferation. Leibiger et al. reported that IR-B promotes glucokinase gene transcription via PI3K signaling (43), consistent with our findings. While many studies suggest long-chain ω-3 polyunsaturated fatty acids (PUFAs) confer cancer-protective effects, their role in cancer progression remains debated (44, 45). We identified ω-3 PUFAs as the primary upregulated fatty acids in IR-B knockout mice, alongside increased expression of their receptor GPR120 and lipogenesis-related genes (Figure 7A, E). Wu et al. identified GPR120 as a tumor-promoting receptor in colorectal carcer (46), paralleling our observations in prostate cancer (Figure 7B-D). Clinical studies indicate that prostate cancer patients consuming a high ω-3, low ω-6 diet supplemented with fish oil exhibit reduced Ki-67 indices, reflecting decreased tumor progression and metastasis (47).

Our data show that EPA and DHA, key ω-3 fatty acids, exert protective effects by modulating BCL2/BAX expression and suppressing the PI3K/AKT pathway, thereby reducing prostate cancer progression. Epidemiological evidence supports ω-3 PUFAs’ role in lowering prostate cancer risk (48). PUFAs are precursors for eicosanoids involved in immune modulation and anti-inflammatory responses (48). Patients supplemented with ω-3 and vitamin D have a higher likelihood of reduced prostate-specific antigen (PSA) levels (49). However, some studies report conflicting associations, including positive correlations between high fish oil levels and prostate cancer incidence (50), or links between DHA and higher tumor grades but not low-grade prostate cancer (51). These discrepancies may stem from differences in fatty acid measurements, disease stage, or tumor metastatic status, highlighting the complexity of PUFA effects in prostate cancer.

Further investigation is needed to determine whether IR isoform alterations represent an inherent aspect of prostate cancer progression, particularly in castration-resistant disease. Our study demonstrates that IR-B knockout dysregulates glucose metabolism and induces compensatory increase in ω-3 PUFAs, which promote GPR120 expression and enhance glucose uptake via GLUT1 (Figure 10). This links IR isoform modulation to glucolipid metabolic dysregulation, offering new insights into prostate carcinoma development.

**Figure 10.**
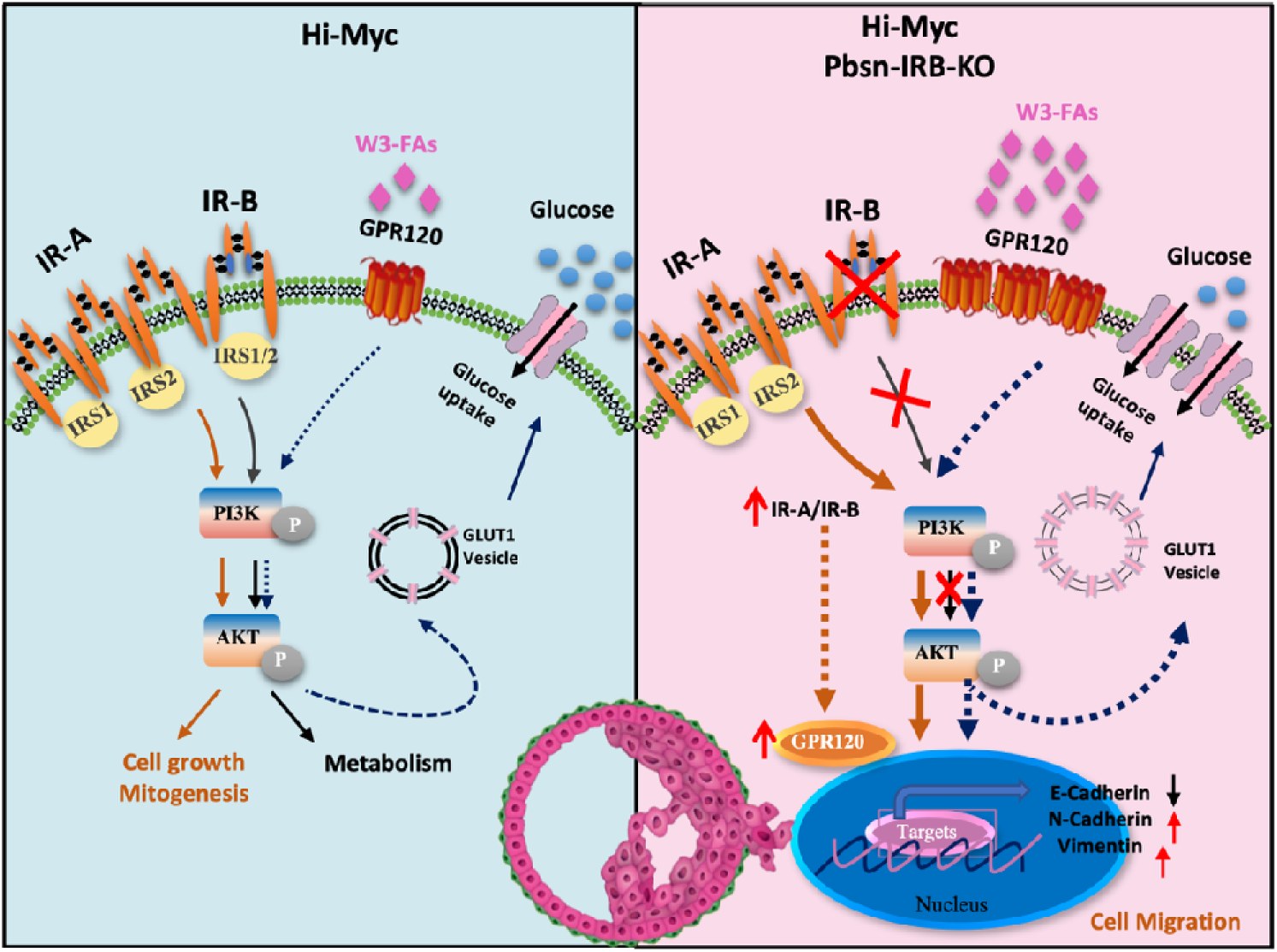
Compensatory regulation of IR-B in glucose and lipid metabolism. Left: Dysregulation of IR isoforms mediates cell proliferation and glucose metabolism via PI3K/AKT signaling in *Hi-Myc* mice. Right: IR-B knockout leads to glucose metabolism disorder and compensatory increase of ω-3 polyunsaturated fatty acids, which upregulate GPR120 expression and enhance glucose uptake through GLUT1.

## MATERIALS AND METHODS

### Animal

IR-B Floxed mice were generated using a Frt-neo-Frt-LoxP conditional targeting vector and embryonic stem (ES) cell aggregation method, as previously described (47). Transgenic Pbsn-Cre mice, which express Cre recombinase under the control of the rat probasin (Pbsn) gene promoter to target prostate epithelial cells, were obtained from The Jackson laboratory. *Hi-Myc* mice, expressing the human *c-MYC* gene under the ARR2Pb promoter specifically in the prostate, were sourced from the NCI Mouse Repository. Td-Tomato reporter mice, carrying a LoxP-flanked STOP cassette upstream of the eGFP fluorescent protein inserted into the Gt (ROSA)26Sor locus, were also purchased from The Jackson laboratory. All mice were housed in polypropylene cages under a standard 12-hour light/dark cycle at 22 ± 2 °C and provided with standard laboratory chow and water ad libitum. All animal procedures were approved by the Dalian Medical University Institutional Animal Care and Use Committee.

### Samples and clinicopathological data

A total of 28 surgically resected prostate cancer specimens and adjacent normal prostate tissue were collected from the Second Hospital of Dalian Medical University between January 2017 and January 2020. None of the patients received any preoperative treatment. Histologic diagnosis was confirmed independently by two expert pathologists. The use of clinical specimens complied with the Declaration of Helsinki and was approved by the Research Ethical Committee of Dalian Medical University.

### Mouse prostate microdissection and histopathologic analysis

The anterior, ventral, and dorsolateral lobes of mouse prostates were carefully dissected under a microscope to remove surrounding adipose and connective tissues. The tissues were then fixed in 10% formalin and processed using standard histological techniques.

### Isolation of RNA and reverse transcriptase PCR quantification

Total RNA was extracted from tissues using TRIzol reagent (Takara) following the manufacturer’s protocol. Complementary DNA (cDNA) was synthesized from total RNA or purified small RNAs using the TransScript One-Step gDNA Removal and cDNA Synthesis SuperMix (Transgen Biotech) according to the manufacturer’s instructions. PCR amplification conditions were as follows: initial denaturation at 94°C for 3 min; 35 cycles of denaturation at 94°C for 30s, annealing at 55°C for 45s, and extension at 72°C for 1min; followed by a final extension at 72°C for 5 min. Real-time quantitative PCR (qPCR) was performed using TransStart Tip Green qPCR SuperMix (Transgen Biotech) on an ABI 7900HT FAST Real-time PCR System (Applied Biosystems, USA). Primer sequences used for qPCR are listed in Supplementary Table 1.

### Western immunoblotting

Western blotting was performed as previously described (9). The following primary antibodies were used: IRS, phospho-IRS1(Tyr896), GSK3β and phospho-GSK3β(Ser9), GPR120, GLUT1, Bcl-2, BAX (Abbkine, USA); IR and phospho-IR, AR, AKT1 and phospho-AKT, PI3K p85 alpha, GAPDH (Proteintech, Wuhan, China); Ki-67 (Cell Signaling Technology, USA); GLUT12 (Boster Bioengineering Co., Ltd., Wuhan); GSK3α/β and phospho-GSK3β (Y216) and phosphor-GSK3α (Y279) (Abcam,USA). Protein bands were visualized using the SuperLumia ECL HRP Substrate Kit (Abbkine, USA). Quantitative analysis was conducted with Quantity One software (Bio-Rad, Hercules, CA, USA).

### Immunohistochemistry

Immunohistochemistry (IHC) was performed on formalin-fixed, paraffin-embedded primary prostate tumor tissues and adjacent normal prostate tissues, which were histopathologically and clinically diagnosed at the Department of Breast Oncology, Second Hospital of Dalian Medical University. Tissue sections were deparaffinized, rehydrated, and subjected to antigen retrieval by microwave treatment in 10 mM citrate buffer (pH 6.0) for 10 minutes. Endogenous peroxidase activity was blocked with 0.3% hydrogen peroxide for 15 minutes. Sections were incubated overnight at 4°C with the following primary antibodies: anti-human and anti-mouse INSR (1:100 dilution), anti-mouse Ki-67 (1:100 dilution), anti-mouse AR (1:100 dilution). Subsequently, sections were incubated with HRP-conjugated secondary antibodies for 30 minutes and developed using 3’-3’ diaminobenzidine (DAB) as the chromogen substrate.

### Analysis of gene expression from databases

Gene expression data were obtained from the ONCOMINE database (Magee Prostate Statistics 2001, Tomlins Prostate Statistics 2007), GEPIA (Gene Expression Profiling Interactive Analysis) database, and The Cancer Genome Atlas (TCGA). Data analyses and figure generation were performed using GraphPad Prism software. In dot plot graphs, each dot represents an individual sample, with results presented as the median and interquartile range.

### Statistical analysis

All data were derived from at least three independent experiments. Statistical analyses were performed using GraphPad Prism version 7.0 (GraphPad Software, San Diego, CA, USA. Differences between two groups evaluated using the two-tailed Student’s t-test for independent samples. Data are presented as mean ± SEM. A p-value of less than 0.05 was considered statistically significant. Statistical significance is indicated as follows: \**P* < 0.05; \*\**P* < 0.01; \*\*\**P* < 0.001.

## LIST OF ABBREVIATIONS

INSR: Insulin Receptor
TCGA: The Cancer Genome Atlas
IHC: immunohistochemistry
IR-A: Insulin Receptor Isoform A
IR-B: Insulin Receptor Isoform B
GPR120: G protein-coupled receptor
GLUT1: Glucose Transporter 1
IGF-I: Insulin-Like Growth Factor I
IGF-II: Insulin-Like Growth Factor II
PUFA: Polyunsaturated fatty acids
ω-3: omega-3 fatty acids
ω-6: omega-6 fatty acids

## ACKNOWLEDGMENTS

We are grateful to all patients for enrolling in this study. This study was funded by the Ministry of Science and Technology (“National Key R&D Program of China” No. 2021YFA0805100, 2022YFE0132200) and National Natural Science Foundation of China (NSFC) No. 82370866 to Y. Wu, No.81872156 to M. Li, and NO.82103132 and Natural Science Foundation of Liaoning Province (NO. 2019-MS-098) to G. Huang.

## DECLARATIONS

### Ethics approval and consent to participate

All patients provided informed consent prior to participation. The study was approved by the Institutional Review Boards of the Second Hospital of Dalian Medical University and was conducted in accordance with the Declaration of Helsinki.

### Authors approval

All authors have seen and approved the manuscript, and that it hasn’t been accepted or published elsewhere.

### Consent for publication

Not applicable

### Availability of data and Materials

The gene expression data are available from cBioportal database, https://www.oncomine.org/, GEPIA (Gene Expression Profiling Interactive Analysis).

## AUTHOR’S CONTRIBUTIONS

Gena Huang collected, analyzed and interpreted data, wrote the manuscript and performed mouse feeding and dissection. Athba AlQahtani, Jing Huang assisted with data collection, and manuscript revision. Jinyu Li contributed to bioinformatics analysis and data interpretation. Kui Jiang facilitated the collection of clinical prostate samples and related data. Sichen Liu assisted with genotyping, data collection and analysis. Zimeng Song supported data collection and analysis. Yue Xi, and Shujing Wang contributed to manuscript review and revision, and provided some materials and advice throughout the study. Man Li designed and helped develop the methodology, supervised the study, and secured funding. Yingjie Wu led the study’s design and methodology development, provided primary supervision, and secured funding.

## Figure Legends

**Table S1.**
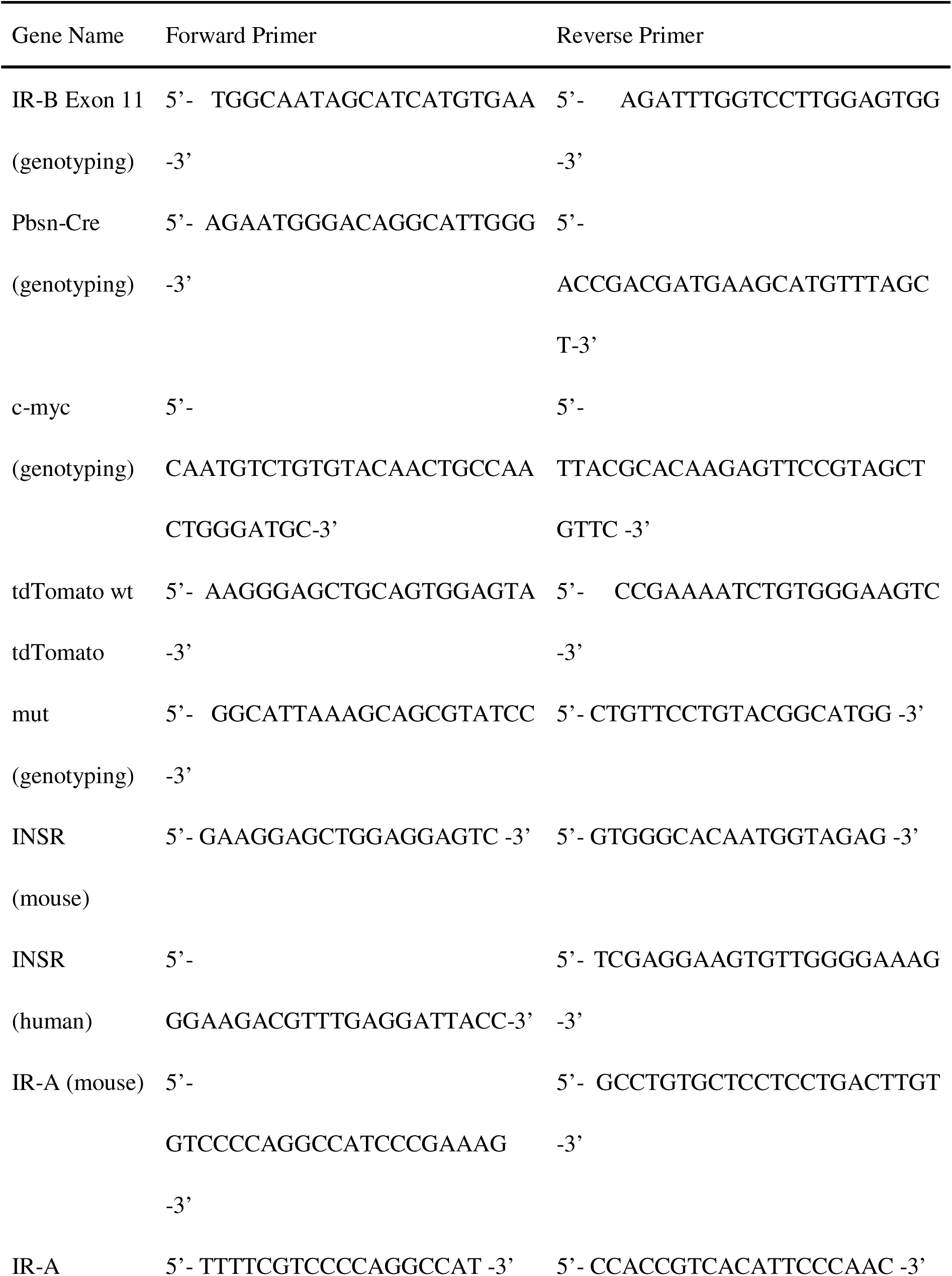
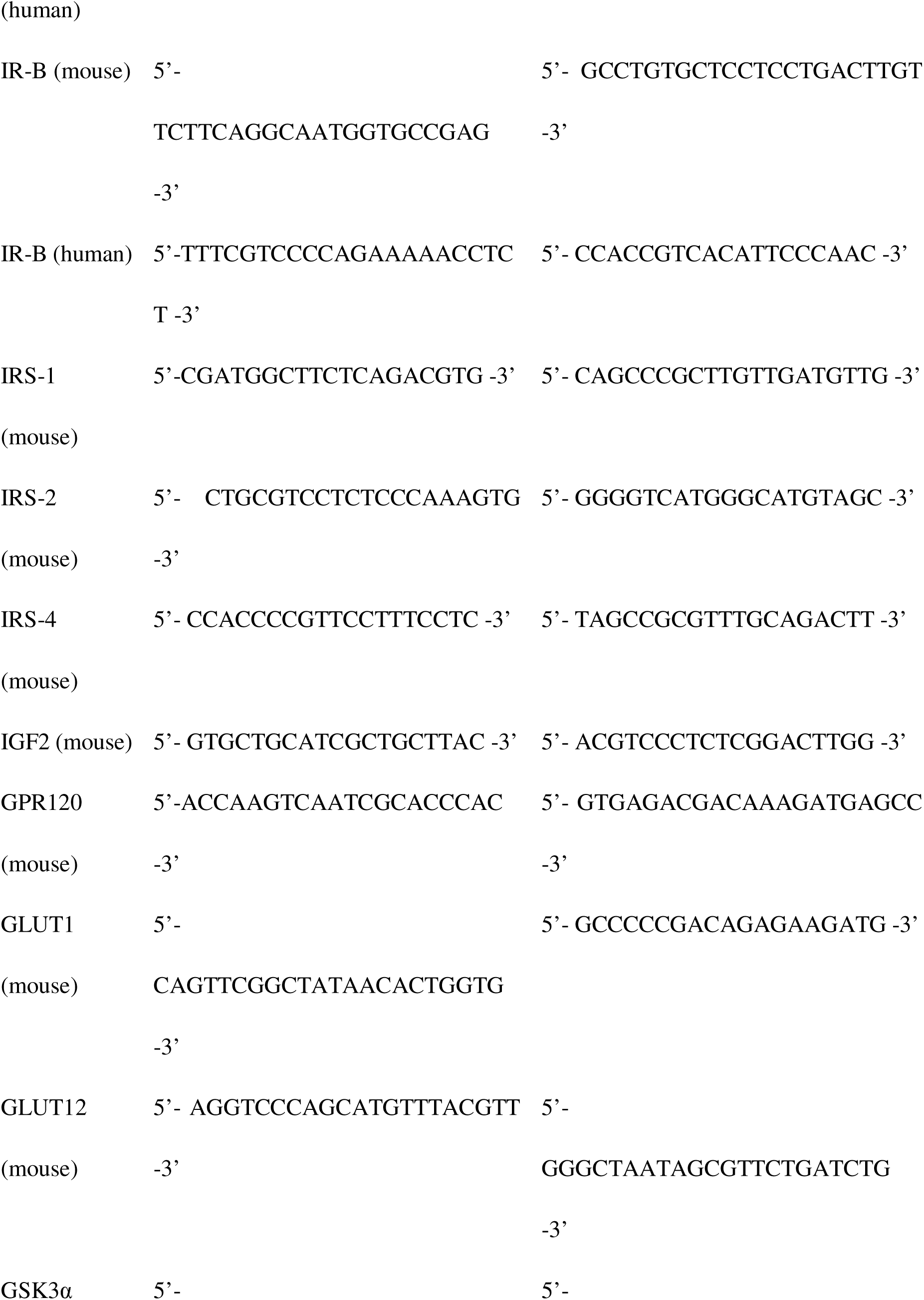
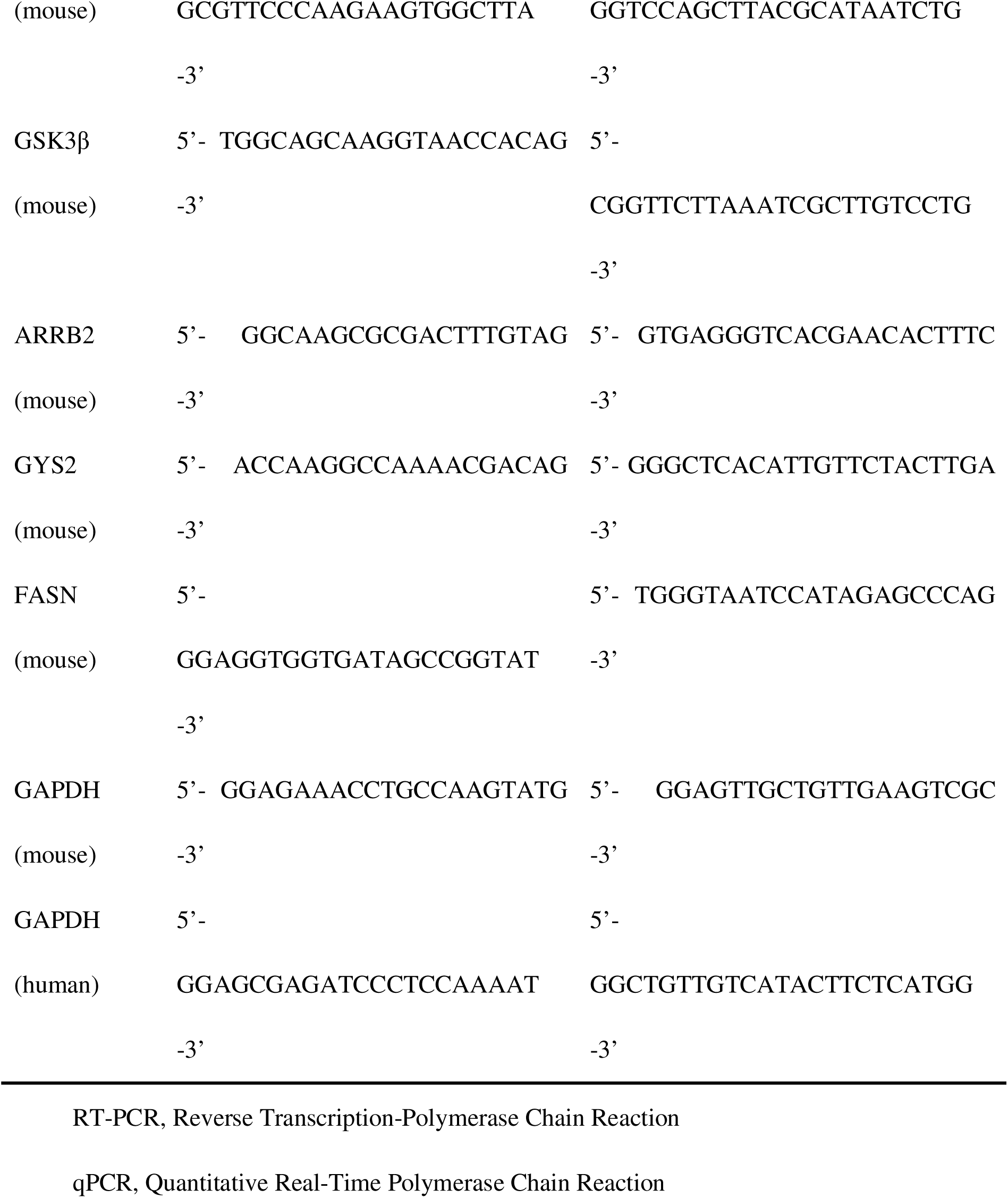
Primer Sequences for RT-PCR and qPCR.

**Table S2.** Enrichment analysis of up-regulated genes by KEGG pathway.

| Biological signaling pathway | *P* value |
| --- | --- |
| Protein processing in endoplasmic reticulum | 6.03E-08 |
| mTOR signaling pathway | 0.000981855 |
| Human papillomavirus infection | 0.070731572 |
| Prostate cancer | 0.001744339 |
| Lipid and atherosclerosis | 0.064247481 |
| Signaling pathways regulating pluripotency of stem cells | 0.040246339 |
| Hippo signaling pathway | 0.063644969 |
| Thyroid hormone synthesis | 0.016932757 |
| TGF-beta signaling pathway | 0.03512445 |
| Serotonergic synapse | 0.061429482 |
| Glycosaminoglycan biosynthesis | 0.005095773 |
| Regulation of lipolysis in adipocytes | 0.026306827 |
| Long-term potentiation | 0.057742982 |
| Ferroptosis | 0.016422633 |
| Endometrial cancer | 0.040468843 |
| Basal cell carcinoma | 0.049692833 |
| Amphetamine addiction | 0.061984883 |
| One carbon pool by folate | 0.02781131 |

**Table S3.**
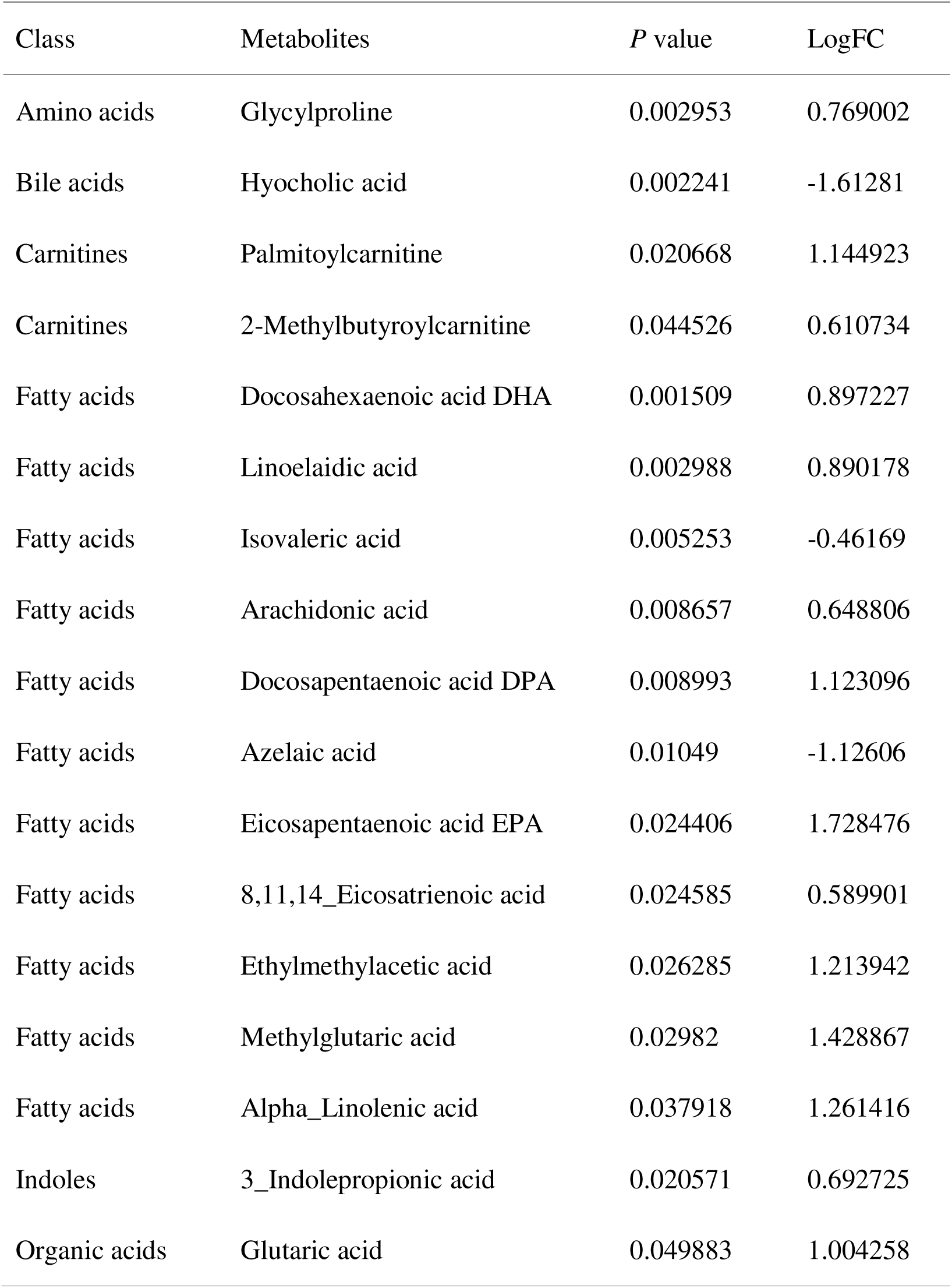
Differential metabolites between IR-B^flox/flox^-Hi-Myc (LL^H^) mice and IR-B^flox/flox^(+)-Hi-Myc (pLL^+H^) mice.

| Class | Metabolites | *P* value | LogFC |
| --- | --- | --- | --- |
| Amino acids | Glycylproline | 0.002953 | 0.769002 |
| Bile acids | Hyocholic acid | 0.002241 | -1.61281 |
| Carnitines | Palmitoylcarnitine | 0.020668 | 1.144923 |
| Carnitines | 2-Methylbutyroylcarnitine | 0.044526 | 0.610734 |
| Fatty acids | Docosahexaenoic acid DHA | 0.001509 | 0.897227 |
| Fatty acids | Linoelaidic acid | 0.002988 | 0.890178 |
| Fatty acids | Isovaleric acid | 0.005253 | -0.46169 |
| Fatty acids | Arachidonic acid | 0.008657 | 0.648806 |
| Fatty acids | Docosapentaenoic acid DPA | 0.008993 | 1.123096 |
| Fatty acids | Azelaic acid | 0.01049 | -1.12606 |
| Fatty acids | Eicosapentaenoic acid EPA | 0.024406 | 1.728476 |
| Fatty acids | 8,11,14_Eicosatrienoic acid | 0.024585 | 0.589901 |
| Fatty acids | Ethylmethylacetic acid | 0.026285 | 1.213942 |
| Fatty acids | Methylglutaric acid | 0.02982 | 1.428867 |
| Fatty acids | Alpha_Linolenic acid | 0.037918 | 1.261416 |
| Indoles | 3_Indolepropionic acid | 0.020571 | 0.692725 |
| Organic acids | Glutaric acid | 0.049883 | 1.004258 |

**Table S4.** Metabolic pathway of differential metabolites.

| Name | *P* value | -LOG(p) | Holm adjust | FDR |
| --- | --- | --- | --- | --- |
| Biosynthesis of unsaturated fatty acid | 1.36E-06 | 13.509 | 0.00011409 | 0.000114 |
| Alpha-Linolenic acid metabolism | 0.083284 | 2.4855 | 1 | 1 |

**Fig. S1.**
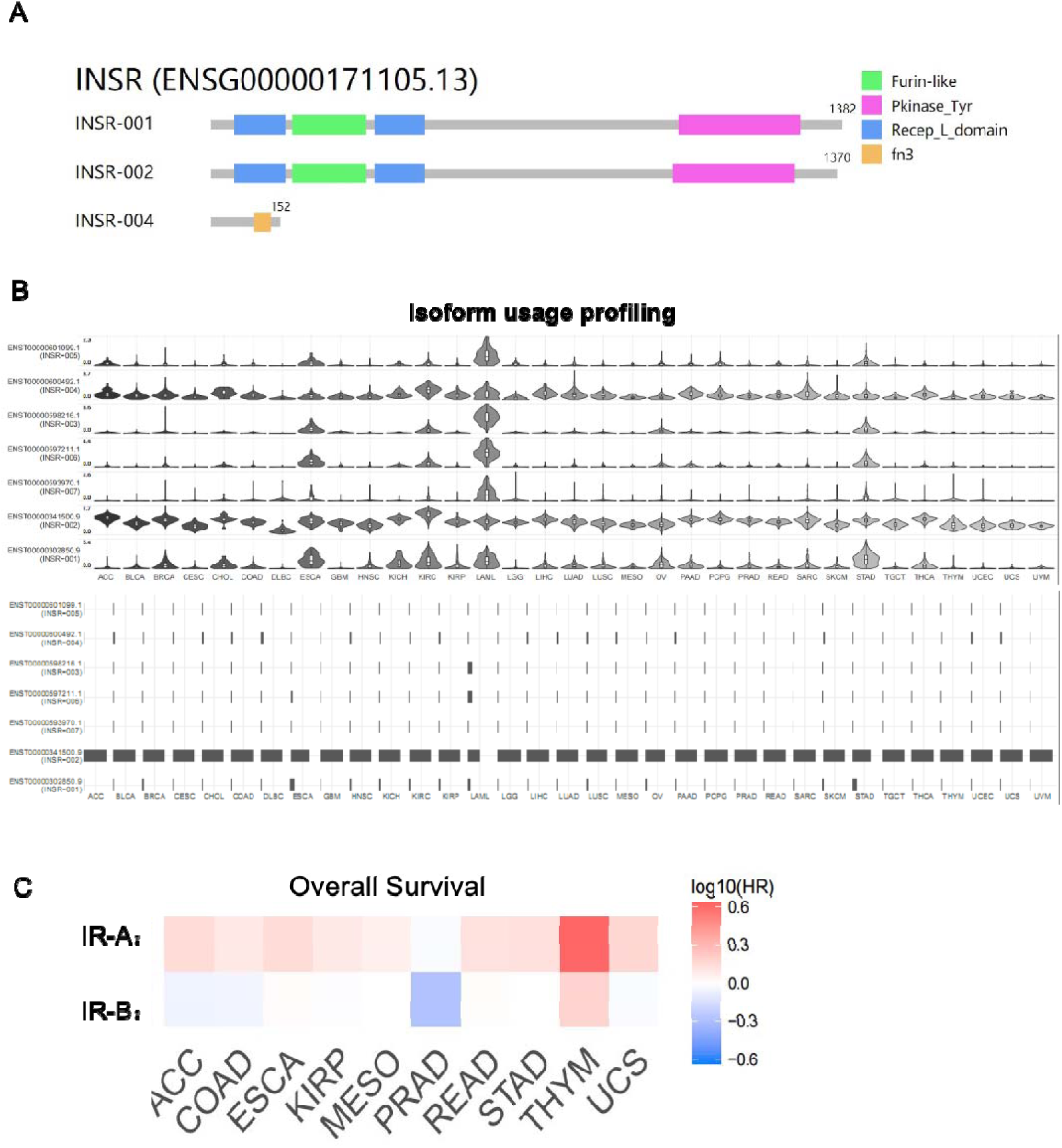

**Fig. S2.**
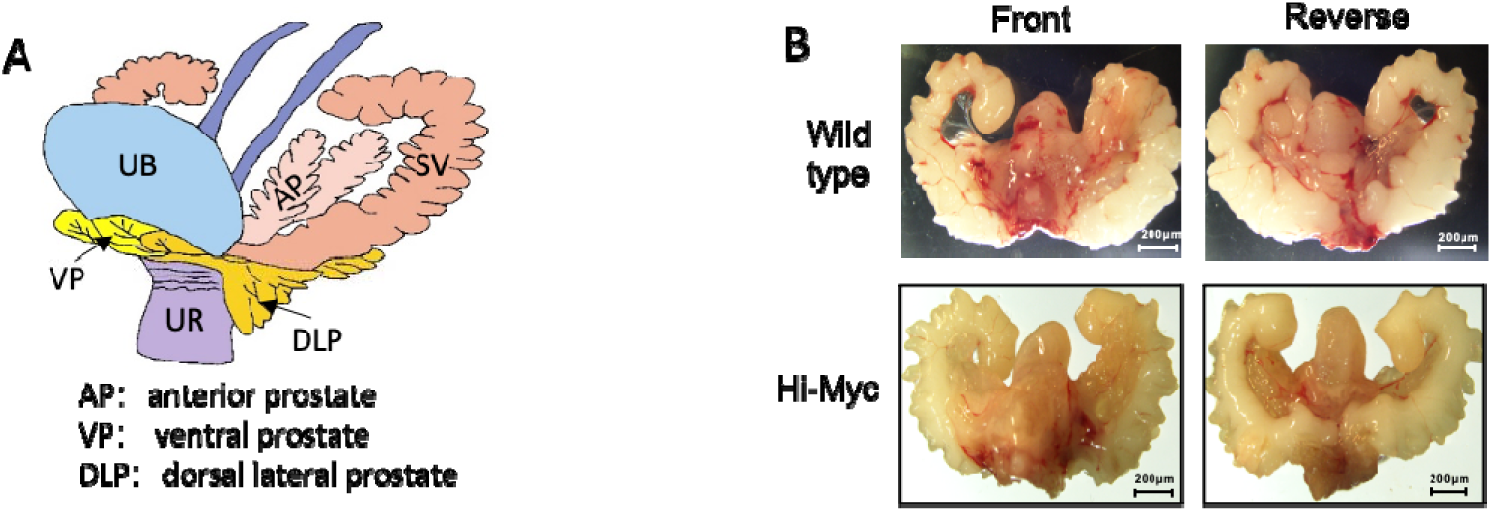

**Fig. S3.**
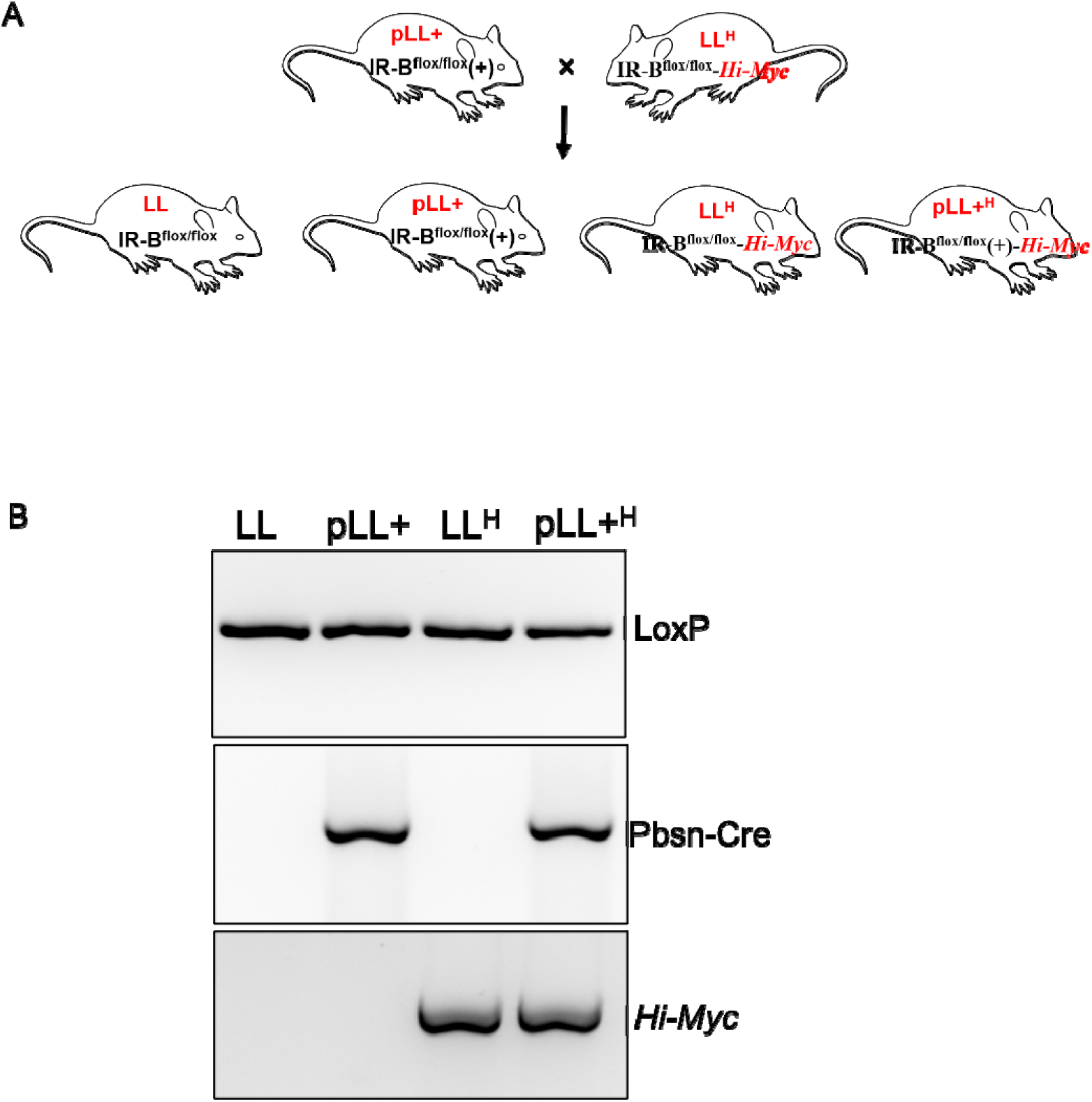

**Fig. S4.**
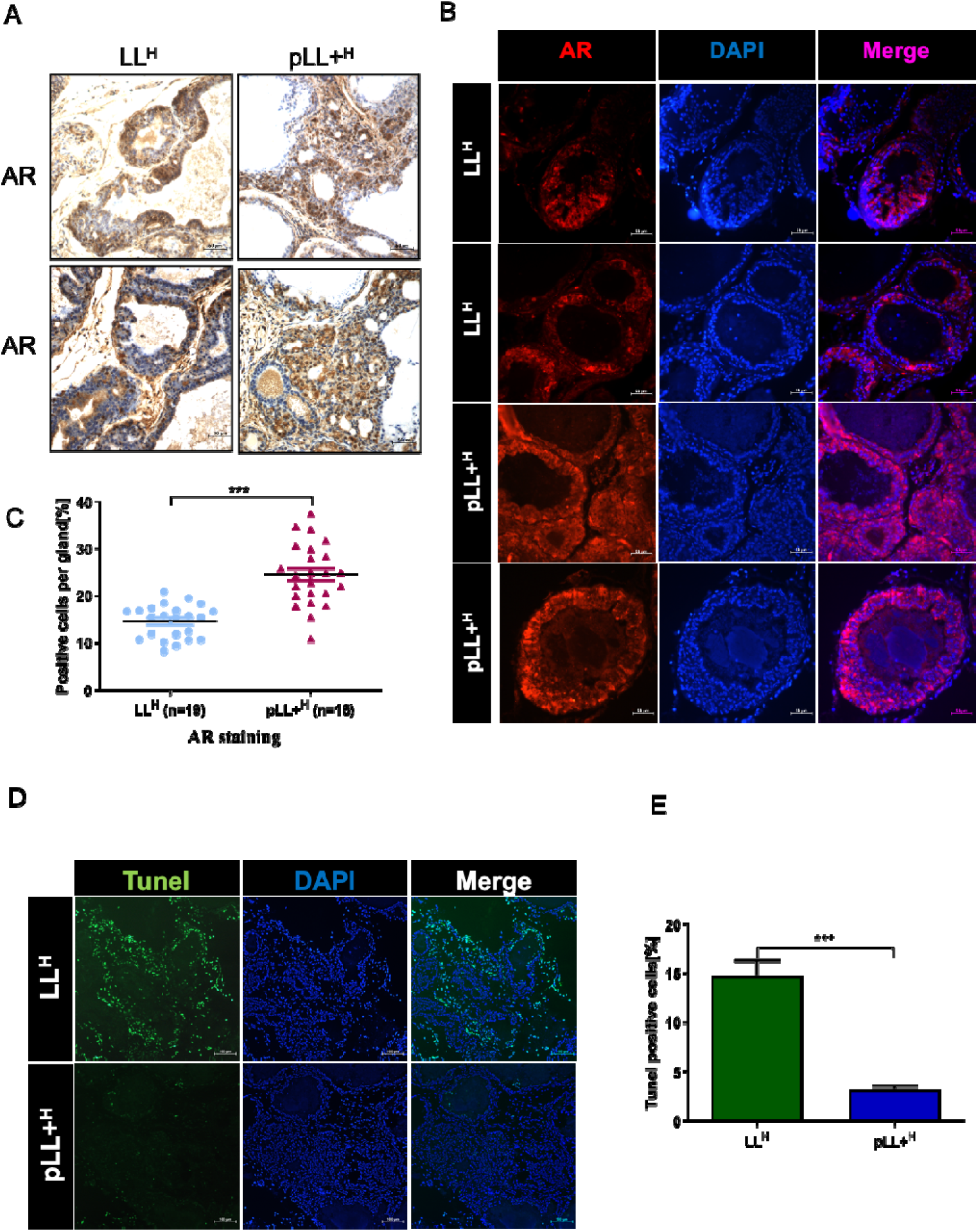

**Fig. S5.**
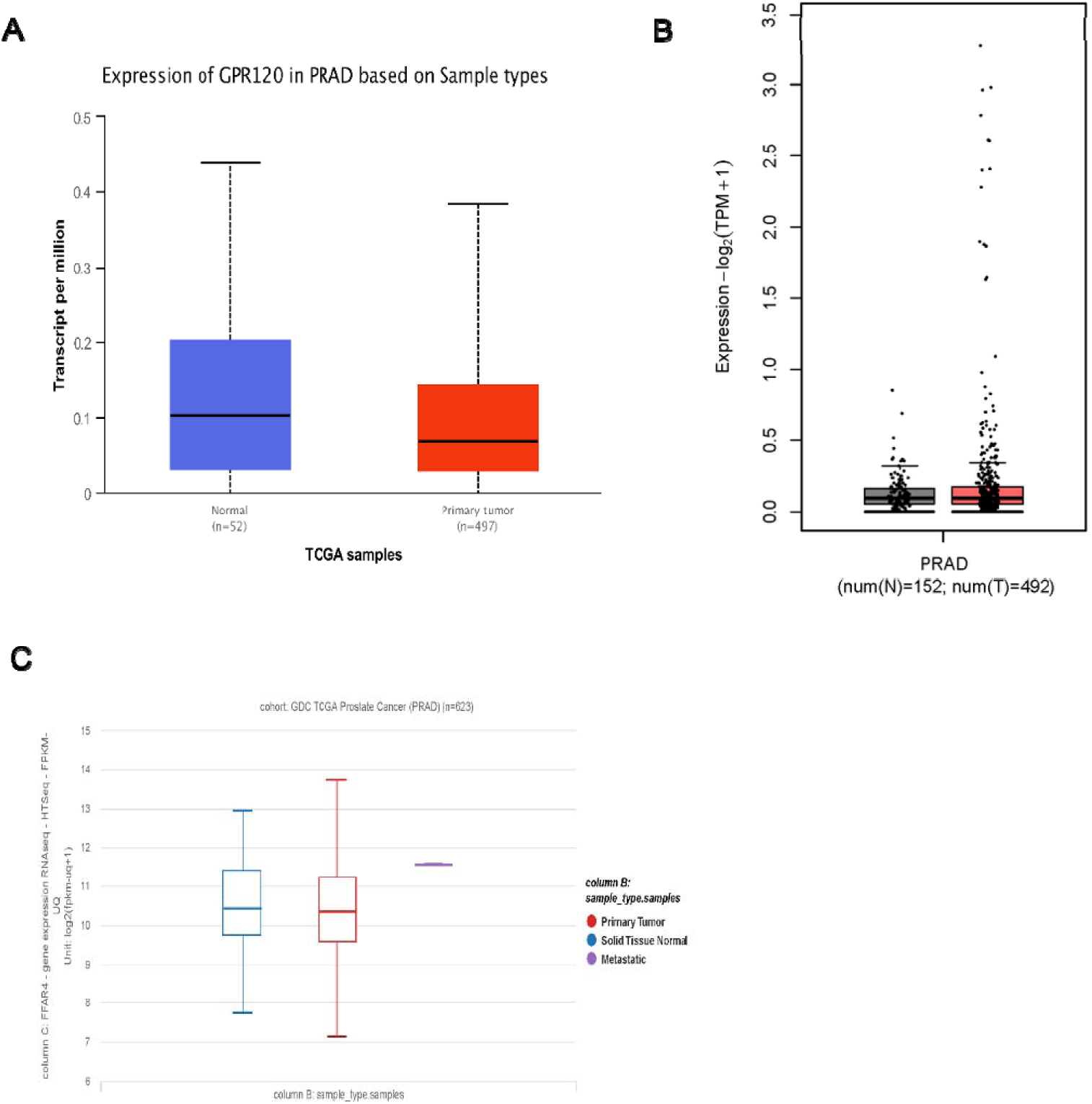

