## Supplemental Figures for "IR-B deficiency and fatty acid dysregulation accelerate prostate cancer progression via PI3K/AKT signaling"

### Slide 1
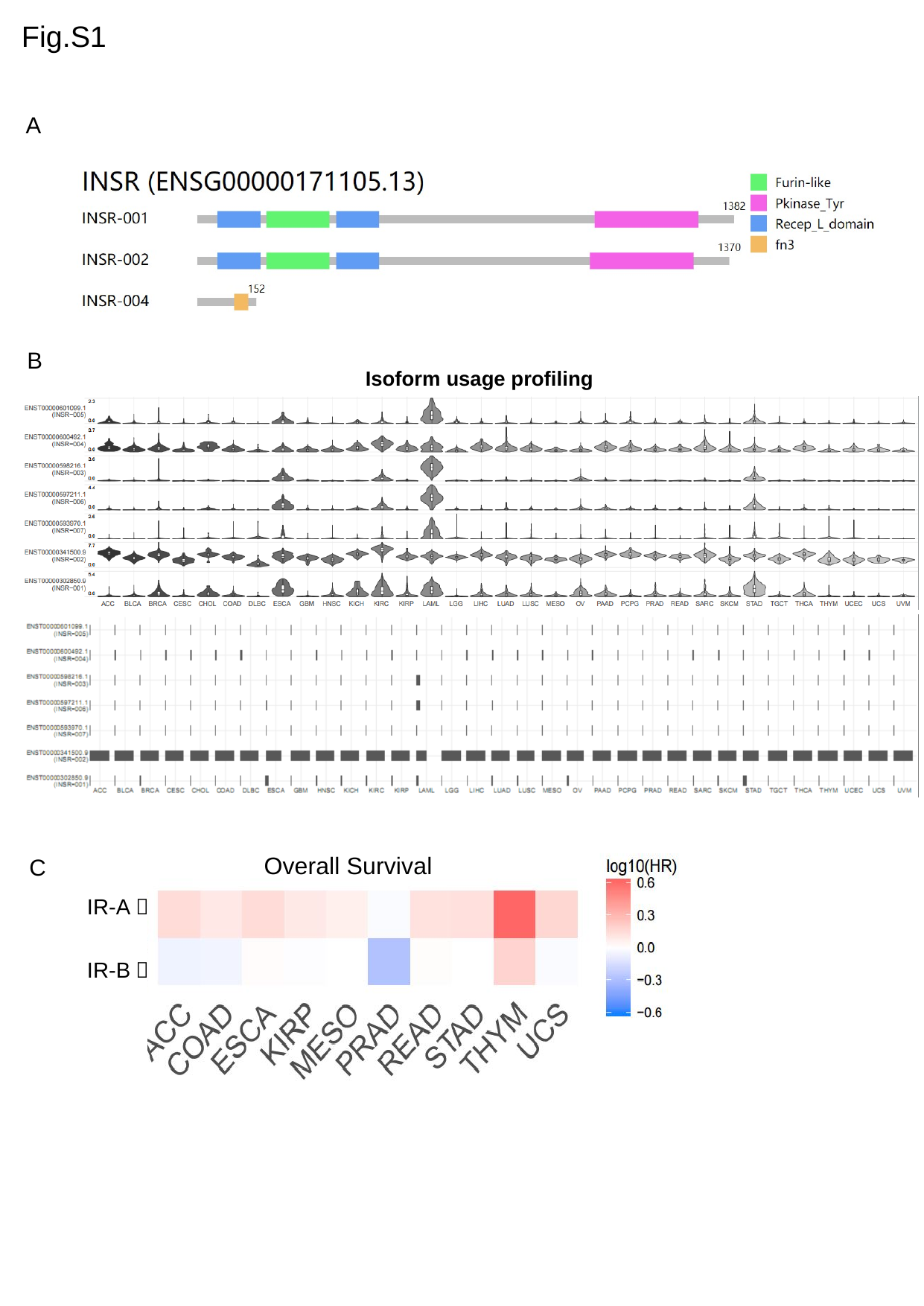

Fig.S1
A
B
Isoform usage profiling
Overall Survival
IR-A：
IR-B：
C

### Slide 2
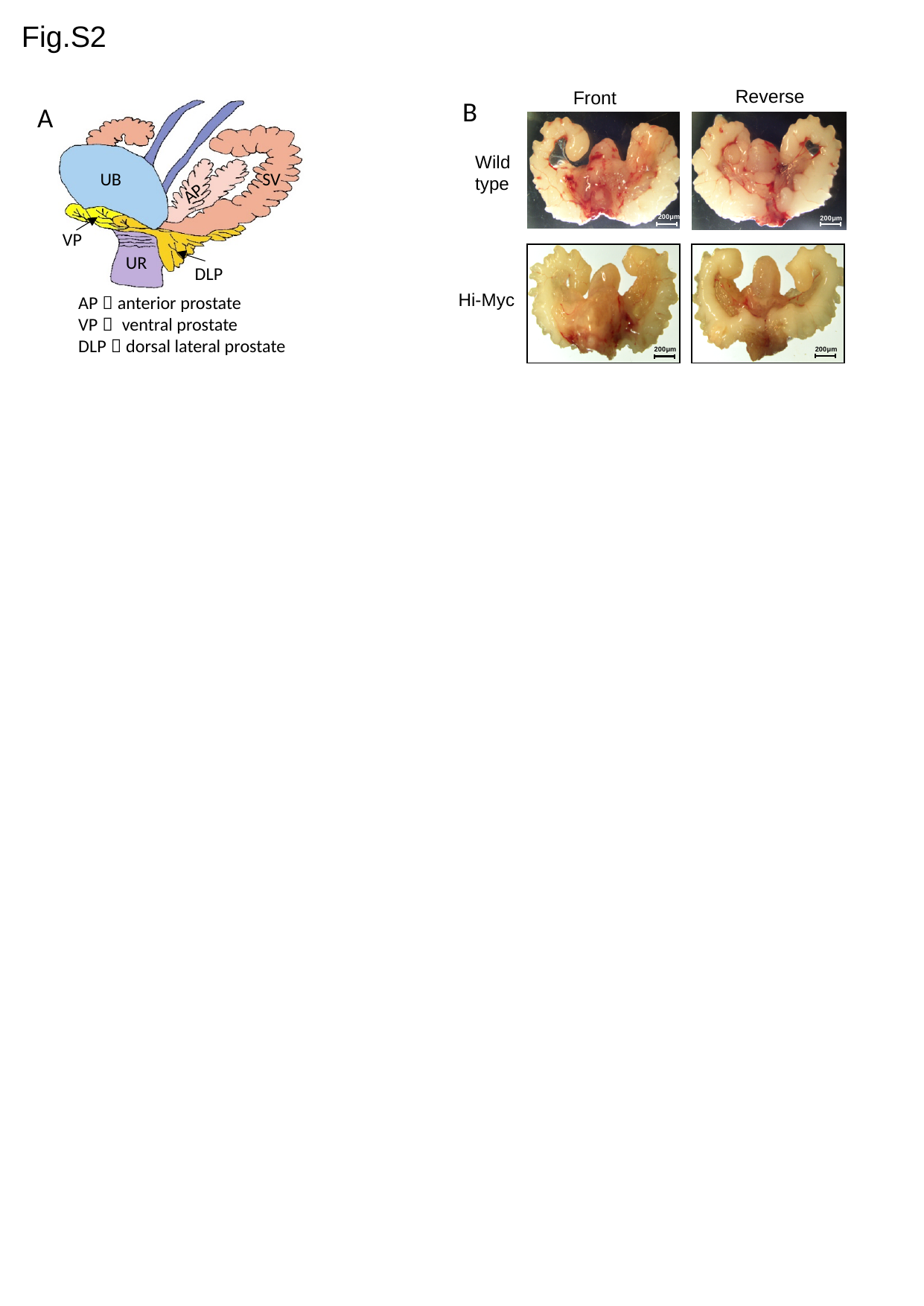

Fig.S2
Reverse
Front
Wild type
200μm
200μm
Hi-Myc
200μm
200μm
B
AP
VP
DLP
AP：anterior prostate
VP： ventral prostate
DLP：dorsal lateral prostate
A
SV
UB
UR

### Slide 3
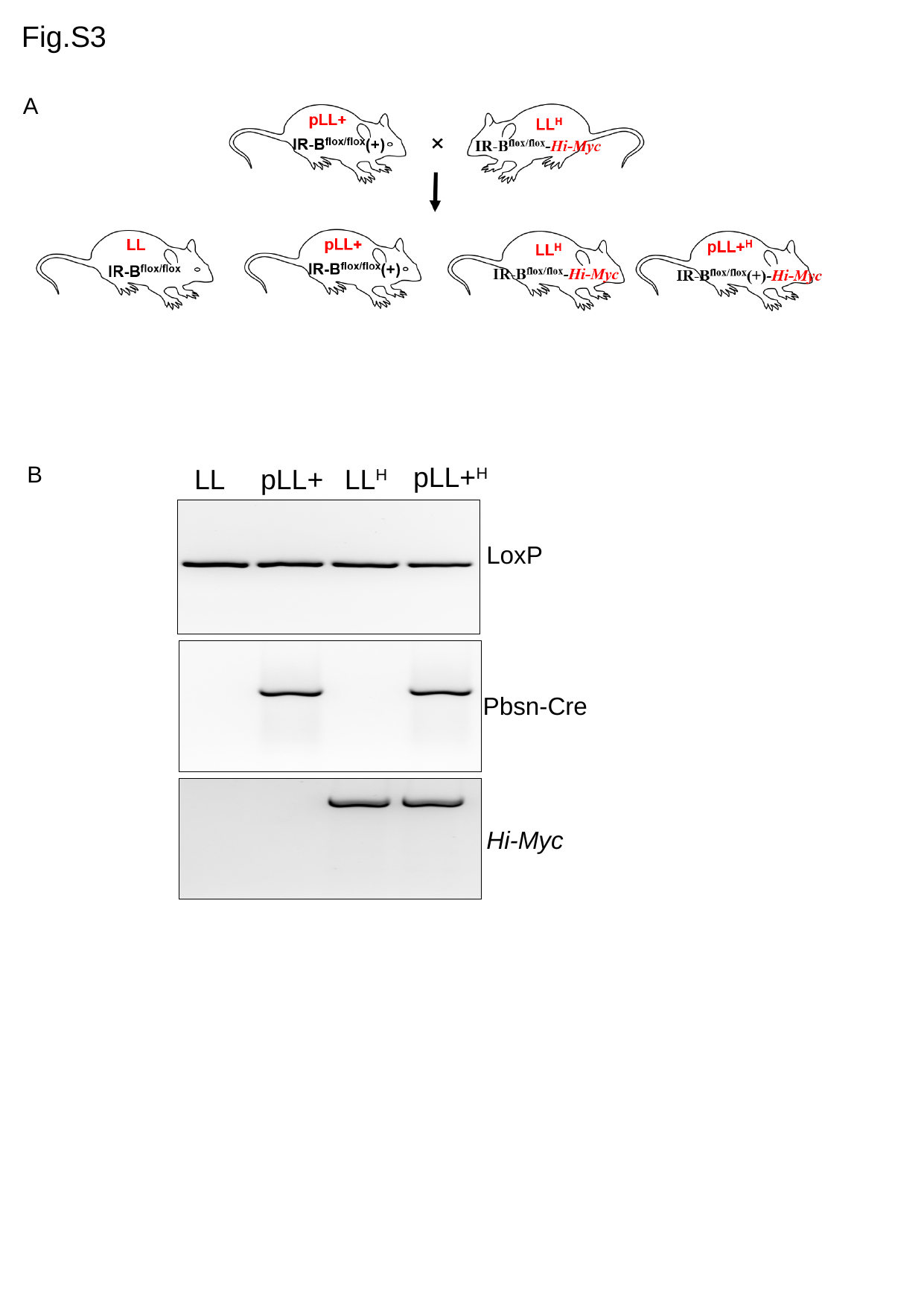

Fig.S3
A
B
pLL+^H^
pLL+
LL^H^
LL
Hi-Myc
LoxP
Pbsn-Cre

### Slide 4
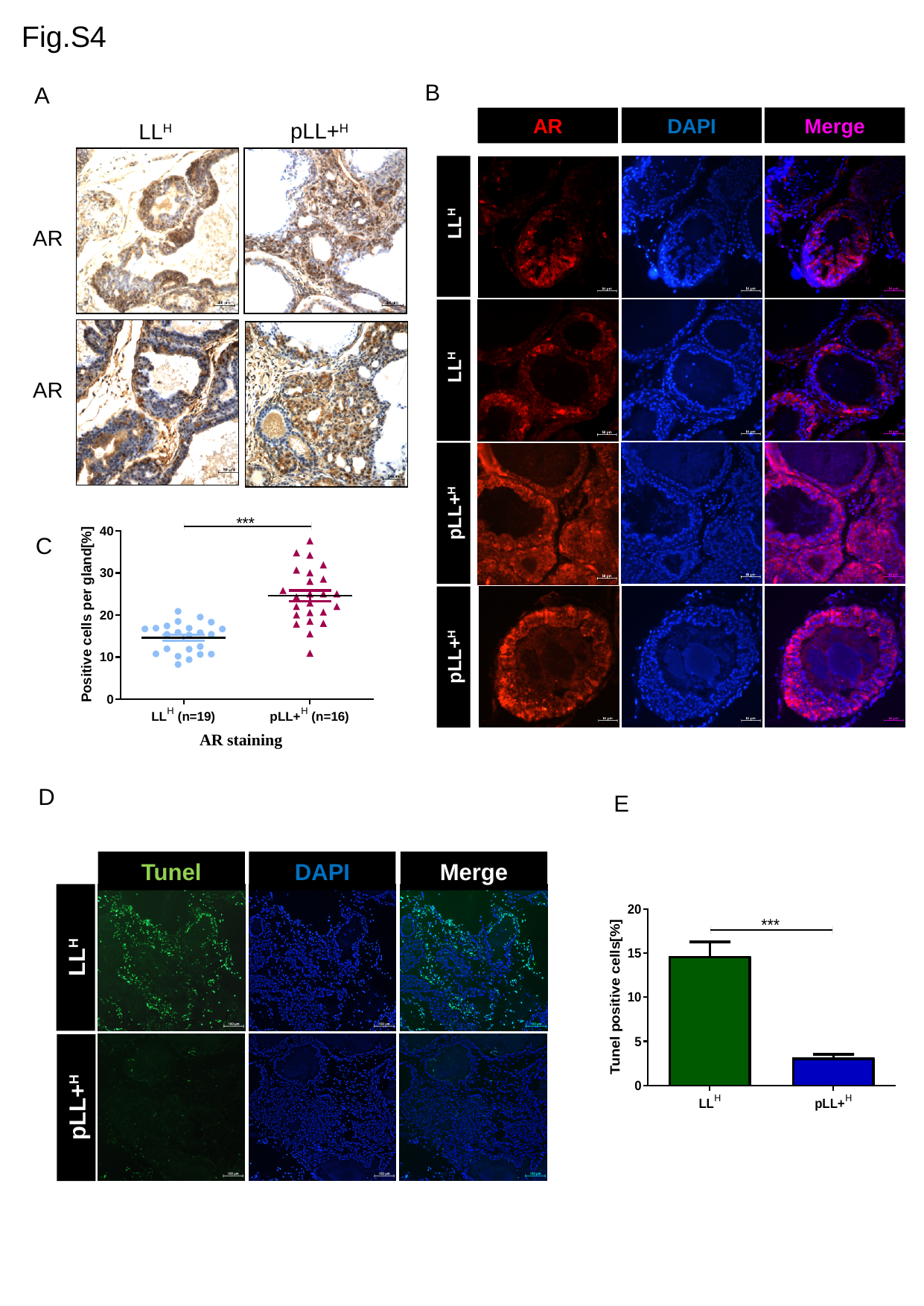

Fig.S4
B
A
DAPI
Merge
AR
 LL^H^
 LL^H^
pLL+^H^
pLL+^H^
pLL+^H^
LL^H^
AR
AR
C
D
E
Tunel
DAPI
Merge
LL^H^
pLL+^H^

### Slide 5
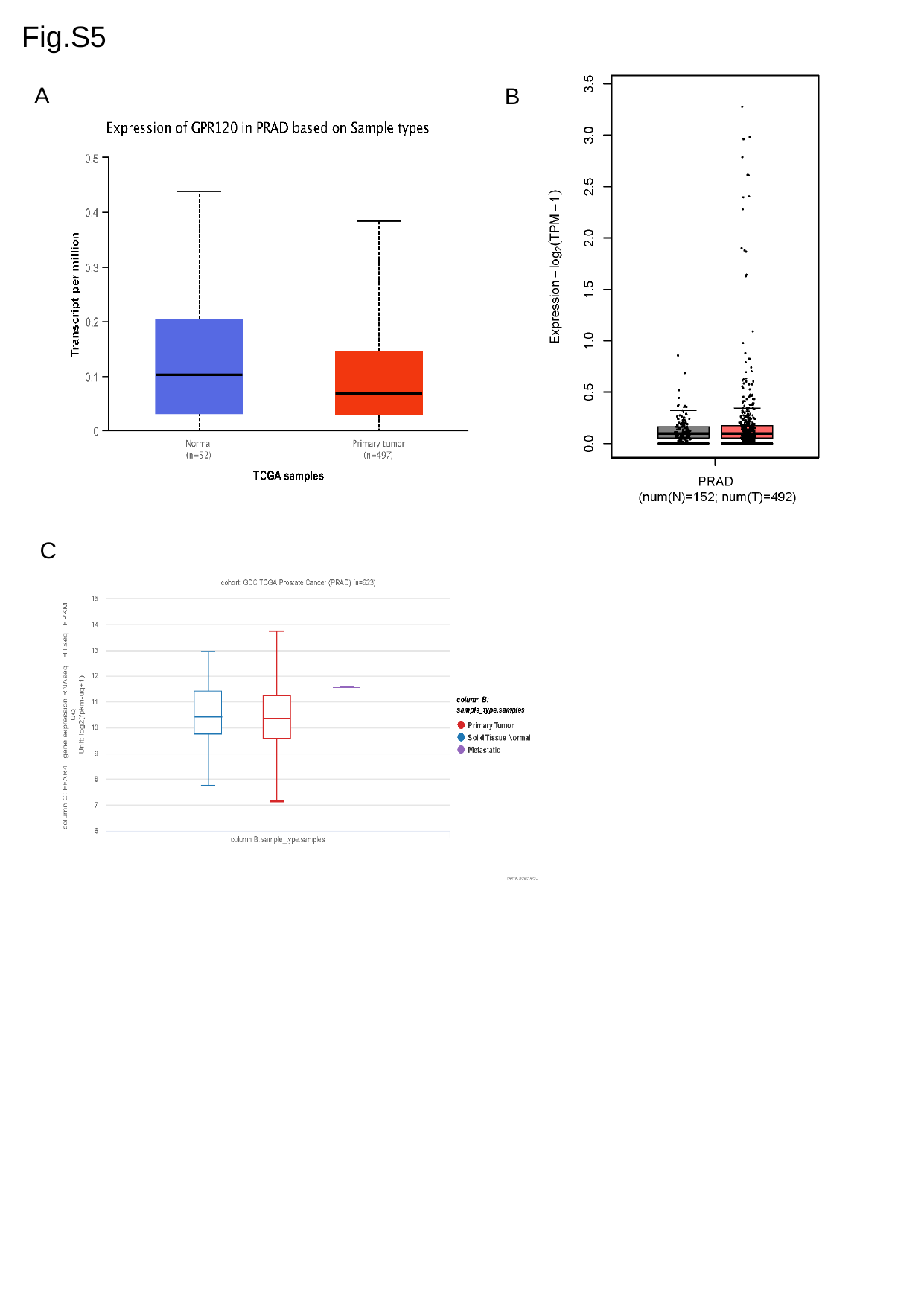

Fig.S5
A
B
C
